# Integrating multiple omics levels using the human protein complexome as a framework, a multi-omics study of inborn errors of metabolism

**DOI:** 10.1101/2023.12.30.573613

**Authors:** Mainak Guharoy, Isabelle Adant, Matthew Bird, Andrea Jáñez Pedrayes, Alexander Botzki, Jonas Dehairs, Stefaan Derveaux, Simon Devos, Geert Goeminne, Francis Impens, Rekin’s Janky, Ruth Maes, Teresa M. Maia, Wouter Meersseman, Daisy Rymen, Johannes V. Swinnen, Delphi Van Haver, Peter Witters, David Cassiman, Bart Ghesquiere

## Abstract

Proteins organize into functional assemblies that drive diverse biological activities. Leveraging a comprehensive dataset of manually curated annotations for the human protein complexome, we investigated biological perturbations at the protein complex level. Using proteomics and transcriptomics data from fibroblasts of patients with inborn errors of metabolism (IEM) and control samples, we globally mapped information onto complex subunits to discern affected processes.

Across the patient cohort (consisting of organic acidaemias, fatty acid oxidation defects and mitochondrial respiratory chain defect IEMs), mitochondrial oxidative phosphorylation emerged as the most perturbed pathway, identified through proteomics datasets. Simultaneously, metabolomics highlighted significant regulation of phospholipids in patients with Fatty Acid and Mitochondrial IEM. Moreover, proteomics analysis also revealed the dysregulation of protein complexes involved in histone (de)acetylation, a finding validated through Western Blot analysis measuring histone acetylation levels. This introduces a novel epigenetic dimension to IEM and metabolic research, suggesting avenues for further exploration.

Our study demonstrates a multiomics data integration concept that maps proteomics and transcriptomics data onto model organism complexomes. This integrative approach can be extended to metabolomics and lipidomics, associating information with complexes having metabolic functions, such as enzymatic complexes. This global strategy for identifying disease-relevant perturbations offers a systems-wide perspective on molecular-level physiological and pathological changes. Such insights are crucial for devising clinical intervention strategies and prioritizing druggable pathways and complexes. The presented methodology provides a foundation for future investigations, emphasizing the importance of integrating multiomics data to comprehensively understand cellular machinery alterations and facilitate targeted therapeutic approaches.

## Introduction

Despite the massive improvements in the field of high-throughput genotyping that enabled the mapping of thousands of genetic variants contributing to disease, the ability to fully grasp the functional consequences of genetic variants on disease necessitates the combination and integration of additional (and complementary) omics fields. In fact, whereas the central dogma of biology insinuates a rather straightforward (linear) relationship between the different omics levels, several studies showed discordances for most transcript-protein pairs^1–4^ evidencing that mRNA levels, on their own, are not always a good predictor of the respective protein abundance. As a solution, multi-omics or the combination of data sets of different omics layers (genomics, transcriptomics, proteomics, lipidomics and metabolomics) was put forward as a powerful approach to understand how disease affects the different cellular processes ranging from transcription to translation and the network of biochemical reactions^5^. The main reason for this relates to the fact that each of the omics levels, on their own, reveals a list of differences that contribute to the disease phenotype and consequently, shed a particular light on the disease mechanisms. For multi-omics, several approaches exist for the integration of the different omics-data levels^6^ of which the most common are correlation or co-mapping^7^.

For a multi-omics study, the field of Inborn Errors of Metabolism (IEM) is of major interest, as the genetic mutation is known to exert its effect across multiple omics levels, from genes, to proteins and metabolites. IEMs are a large group of genetic disorders, which have a variable presentation, incidence and severity^8^. IEMs can present at any time in life, from early development until adulthood, though the majority of IEMs known to date present in the first years of life. These disorders usually occur due to the pathogenic variants in genes essential for various metabolic pathways such as the genes encoding for enzymes, cofactors or transporters. The most common mode of genetic inheritance is autosomal recessive, though X -linked, dominant and mitochondrial inheritance are also described^9,10^. Despite advances in understanding the genetic basis of IEMs, the underlying molecular mechanisms remain largely elusive. This is primarily due to the complexity of the molecular pathways involved in IEMs and the difficulty of accurately measuring and quantifying all the relevant molecules involved in their etiology.

Consequently, applying a multi-omics approach onto different IEMs could provide better insights into the disease mechanisms. In this work, we describe such a multi-omics study investigating the integration of proteomics, transcriptomics and metabolomics datasets of different classes of IEM: organic acidaemias (Cobalamin B and C deficiency), fatty acid oxidation defects (very long-chain acyl-CoA dehydrogenase deficiency and medium-chain acyl-CoA deficiency) and mitochondrial respiratory chain defects (Complex I deficiency). Investigating how a single genetic disorder can affect multiple levels of the biochemical machinery can lead to a better understanding of these disorders, eventually creating possibilities for novel treatment strategies. We investigated to what extent the different levels of -omics could rationalize or even predict the functional consequences of IEMs. We also mapped the correlation between the different omics layers and checked whether multi-omics provided additional advantages for elucidating the pathophysiology of disease. For the data integration, we designed a systems-wide interactomics and complexomics approach based on interrogating manually curated functional and physical protein interaction networks from IntAct^11^ as well as further curated, closely interacting groups of proteins representing well-annotated protein complex assemblies from Complex Portal^12^. This collection of >1200 human protein complexes constitutes the complexome, representing the essential molecular machinery within cells (ranging from the highly stable to those that are dynamically regulated). Therefore, the interaction network together with the annotated complexes provided a functional framework to contextualize the multi-omics data. It enabled us to: 1) query for and observe perturbations in the molecular machinery affected in the IEM cohorts, and 2) map the correspondences across molecular layers. Using this approach, we identified specific protein complexes that are perturbed in IEMs, leading to an enhanced understanding of the aberrant molecular mechanisms, and thereby bringing us closer to explaining the pathophysiology and, ultimately, the phenotype.

## Results

### Overview of the IEM samples and targets identified per omics level

Patient skin-derived fibroblasts from individuals diagnosed with IEM and healthy individuals were processed for the respective omics platforms. The patient cohort was divided into 3 disease categories (**Table 1**): organic acidaemias (OA), fatty acid oxidation defects (FA) and mitochondrial complex I defects (Mito). After performing filtering and pre-processing steps (see Methods), 3140 proteins and 16202 transcripts were retained for the downstream analysis of disease-linked changes. The dataset obtained from the LC-MS based untargeted metabolomics provided a total of 5644 features (positive and negative ionization combined).Mass spectrometry-based proteomics highlights disease-relevant metabolic enzymes

**Table 1.**
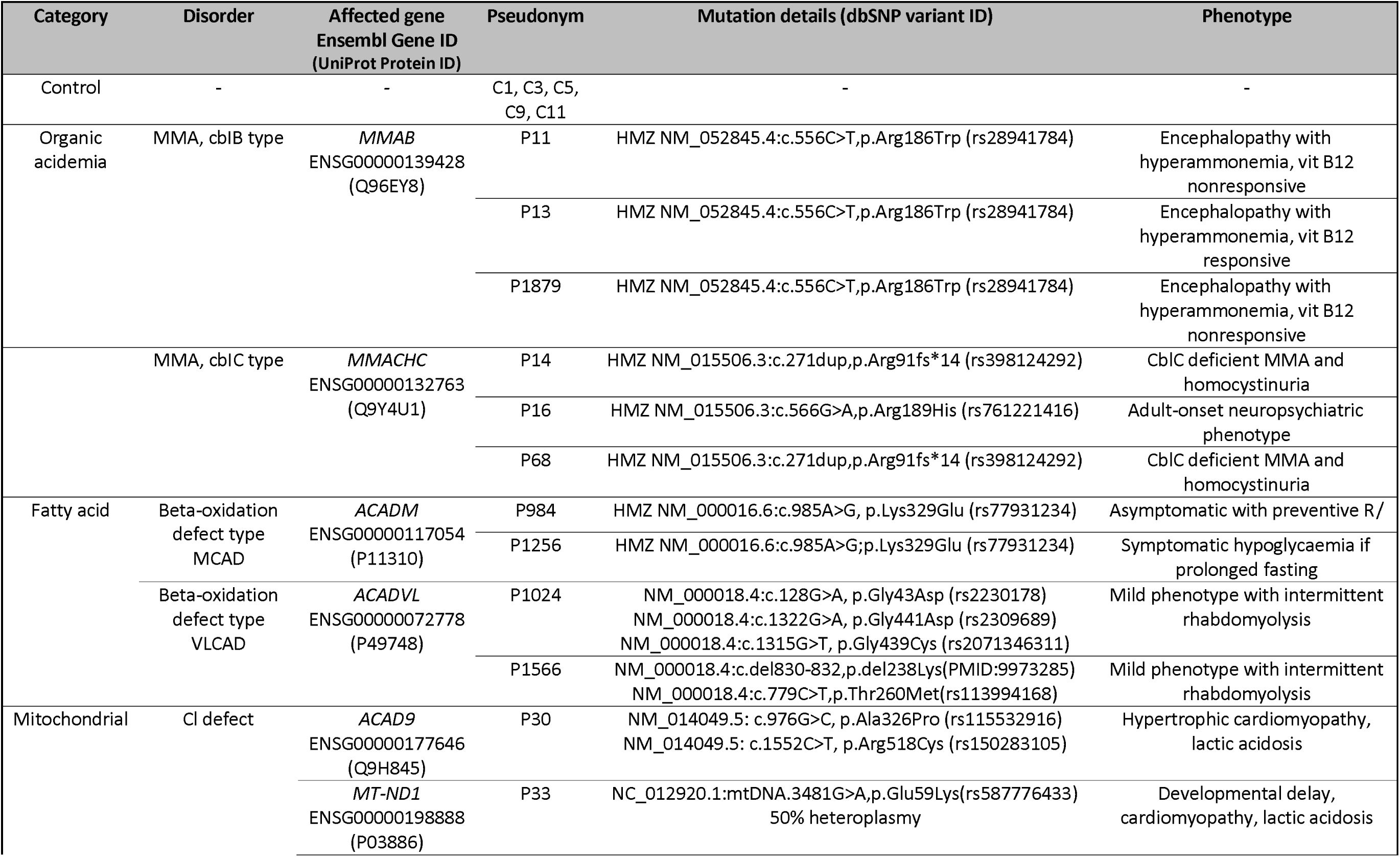
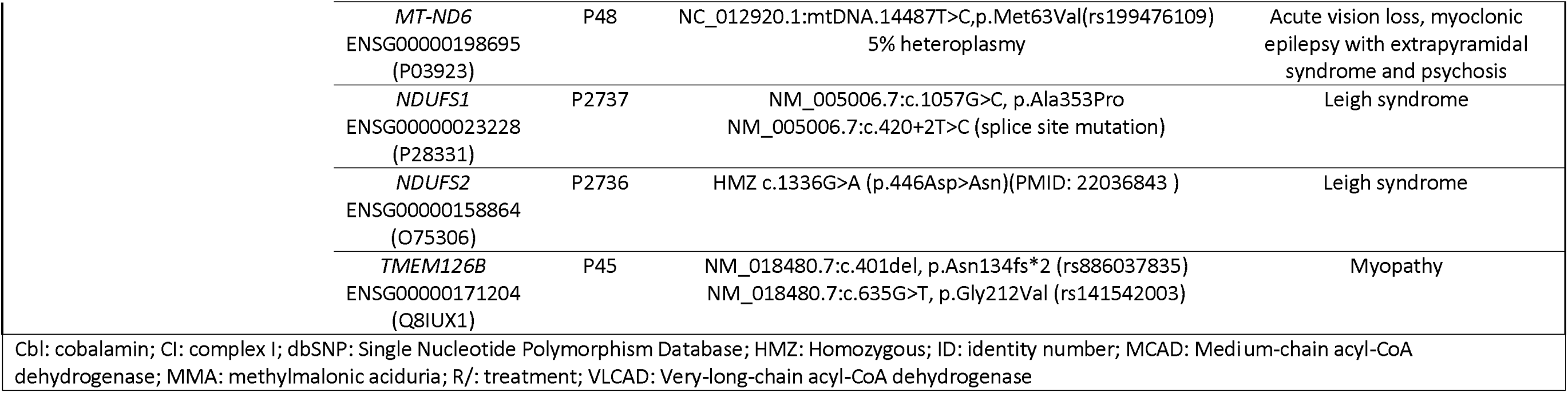
Overview of skin-derived fibroblast cell lines obtained from healthy individuals (controls) and patients with inborn errors of metabolism.

| Category | Disorder | Affected gene<br>Ensembl Gene ID<br>(UniProt Protein ID) | Pseudonym | Mutation details (dbSNP variant ID) | Phenotype |
| --- | --- | --- | --- | --- | --- |
| Control | - | - | C1, C3, C5,<br>C9, C11 | - | - |
| Organic<br>acidemia | MMA, cblB type | *MMAB*<br>ENSG00000139428<br>(Q96EY8) | P11 | HMZ NM_052845.4:c.556C>T,p.Arg186Trp (rs28941784) | Encephalopathy with<br>hyperammonemia, vit B12<br>nonresponsive |
|  |  |  | P13 | HMZ NM_052845.4:c.556C>T,p.Arg186Trp (rs28941784) | Encephalopathy with<br>hyperammonemia, vit B12<br>responsive |
|  |  |  | P1879 | HMZ NM_052845.4:c.556C>T,p.Arg186Trp (rs28941784) | Encephalopathy with<br>hyperammonemia, vit B12<br>nonresponsive |
|  | MMA, cblC type | *MMACHC*<br>ENSG00000132763<br>(Q9Y4U1) | P14 | HMZ NM_015506.3:c.271dup,p.Arg91fs*14 (rs398124292) | CblC deficient MMA and<br>homocystinuria |
|  |  |  | P16 | HMZ NM_015506.3:c.566G>A,p.Arg189His (rs761221416) | Adult-onset neuropsychiatric<br>phenotype |
|  |  |  | P68 | HMZ NM_015506.3:c.271dup,p.Arg91fs*14 (rs398124292) | CblC deficient MMA and<br>homocystinuria |
| Fatty acid | Beta-oxidation<br>defect type<br>MCAD | *ACADM*<br>ENSG00000117054<br>(P11310) | P984 | HMZ NM_000016.6:c.985A>G, p.Lys329Glu (rs77931234) | Asymptomatic with preventive R/ |
|  |  |  | P1256 | HMZ NM_000016.6:c.985A>G;p.Lys329Glu (rs77931234) | Symptomatic hypoglycaemia if<br>prolonged fasting |
|  | Beta-oxidation<br>defect type<br>VLCAD | *ACADVL*<br>ENSG00000072778<br>(P49748) | P1024 | NM_000018.4:c.128G>A, p.Gly43Asp (rs2230178)<br>NM_000018.4:c.1322G>A, p.Gly441Asp (rs2309689)<br>NM_000018.4:c.1315G>T, p.Gly439Cys (rs2071346311) | Mild phenotype with intermittent<br>rhabdomyolysis |
|  |  |  | P1566 | NM_000018.4:c.del830-832,p.del238Lys(PMID:9973285)<br>NM_000018.4:c.779C>T,p.Thr260Met(rs113994168) | Mild phenotype with intermittent<br>rhabdomyolysis |
| Mitochondrial | CI defect | *ACAD9*<br>ENSG00000177646<br>(Q9H845) | P30 | NM_014049.5: c.976G>C, p.Ala326Pro (rs115532916)<br>NM_014049.5: c.1552C>T, p.Arg518Cys (rs150283105) | Hypertrophic cardiomyopathy,<br>lactic acidosis |
|  |  | *MT-ND1*<br>ENSG00000198888<br>(P03886) | P33 | NC_012920.1:mtDNA.3481G>A,p.Glu59Lys(rs587776433)<br>50% heteroplasmy | Developmental delay,<br>cardiomyopathy, lactic acidosis |

|  |  |  |  |
| --- | --- | --- | --- |
| *MT-ND6*<br>ENSG00000198695<br>(P03923) | P48 | NC_012920.1:mtDNA.14487T>C, p.Met63Val(rs199476109)<br>5% heteroplasmy | Acute vision loss, myoclonic<br>epilepsy with extrapyramidal<br>syndrome and psychosis |
| *NDUFS1*<br>ENSG00000023228<br>(P28331) | P2737 | NM_005006.7:c.1057G>C, p.Ala353Pro<br>NM_005006.7:c.420+2T>C (splice site mutation) | Leigh syndrome |
| *NDUFS2*<br>ENSG00000158864<br>(O75306) | P2736 | HMZ c.1336G>A (p.446Asp>Asn)(PMID: 22036843 ) | Leigh syndrome |
| *TMEM126B*<br>ENSG00000171204<br>(Q8IUX1) | P45 | NM_018480.7:c.401del, p.Asn134fs*2 (rs886037835)<br>NM_018480.7:c.635G>T, p.Gly212Val (rs141542003) | Myopathy |
| Cbl: cobalamin; Cl: complex I; dbSNP: Single Nucleotide Polymorphism Database; HMZ: Homozygous; ID: identity number; MCAD: Medium-chain acyl-CoA dehydrogenase; MMA: methylmalonic aciduria; R/: treatment; VLCAD: Very-long-chain acyl-CoA dehydrogenase |  |  |  |

To identify the molecular mechanisms underlying disease-specific perturbations, we performed proteomics analysis on the IEM patient samples. Overall, we quantified a total of 3140 unique proteins over all samples and identified 24, 109 and 100 proteins regulated significantly for the OA, FA and Mito IEM patient groups, respectively, relative to control samples, based on a statistical assessment (FDR <= 0.05) and fold change cutoff of 2 (**Figure 1**). Several of these significantly regulated proteins were metabolic enzymes (**Figure 1**) that are linked to the relevant biological functions of the reported patient phenotypes. For example, BCAT1 is a branched-chain amino acid aminotransferase that participates in the metabolism of the essential branched chain amino acids leucine, isoleucine, and valine. Here, BCAT1 was found to be upregulated in OA IEM samples (**Figure 1A**), which could in turn affect levels of leucine, mTORC1 signaling as well as glycolysis^13^. In FA IEM samples, very long-chain specific acyl-CoA dehydrogenase (ACADVL) was one of the enzymes downregulated (**Figure 1B**). ACADVL is an acyl-CoA dehydrogenase that catalyzes the first step of mitochondrial fatty acid beta-oxidation, breaking fatty acids into ultimately acetyl-CoA, and thereby allowing energy production from fats^14^. Indeed, ACADVL is mutated in two of the four FA IEM samples analyzed in this study (**Table 1**). ACAD11, another acyl-CoA dehydrogenase, was also found to be similarly regulated. Interestingly, several subunits of mitochondrial membrane respiratory chain complexes and proteins involved in their assembly were also found to be affected in FA IEM samples (NDUFAB1, MT-ND5, ATP5L, COX20 and DNAJC11; **Figure 1B**). Similarly, in the case of the Mito IEM patient samples, there was a pronounced effect on several subunits of mitochondrial electron transport chain complexes (**Figure 1C**).

**Figure 1.**
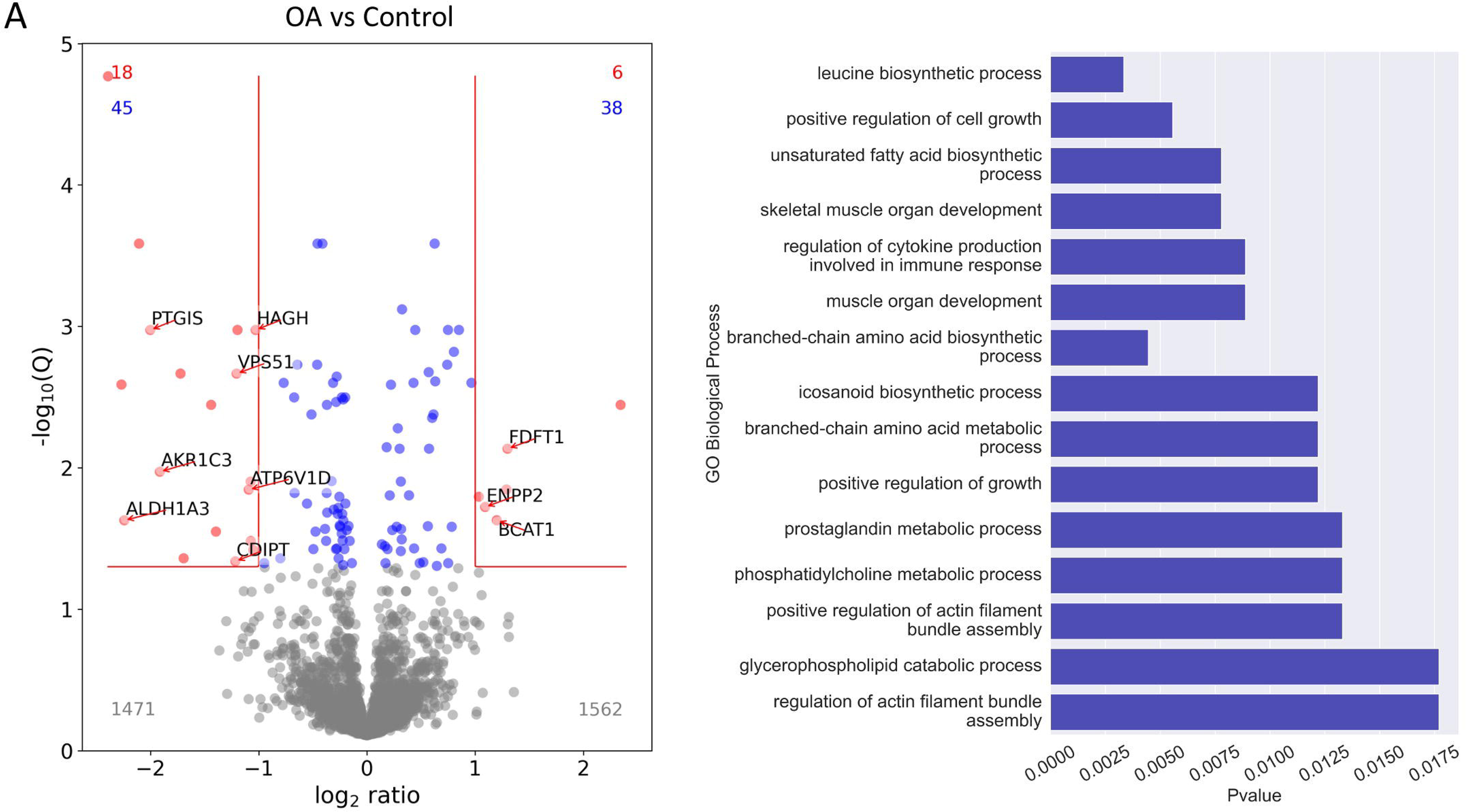
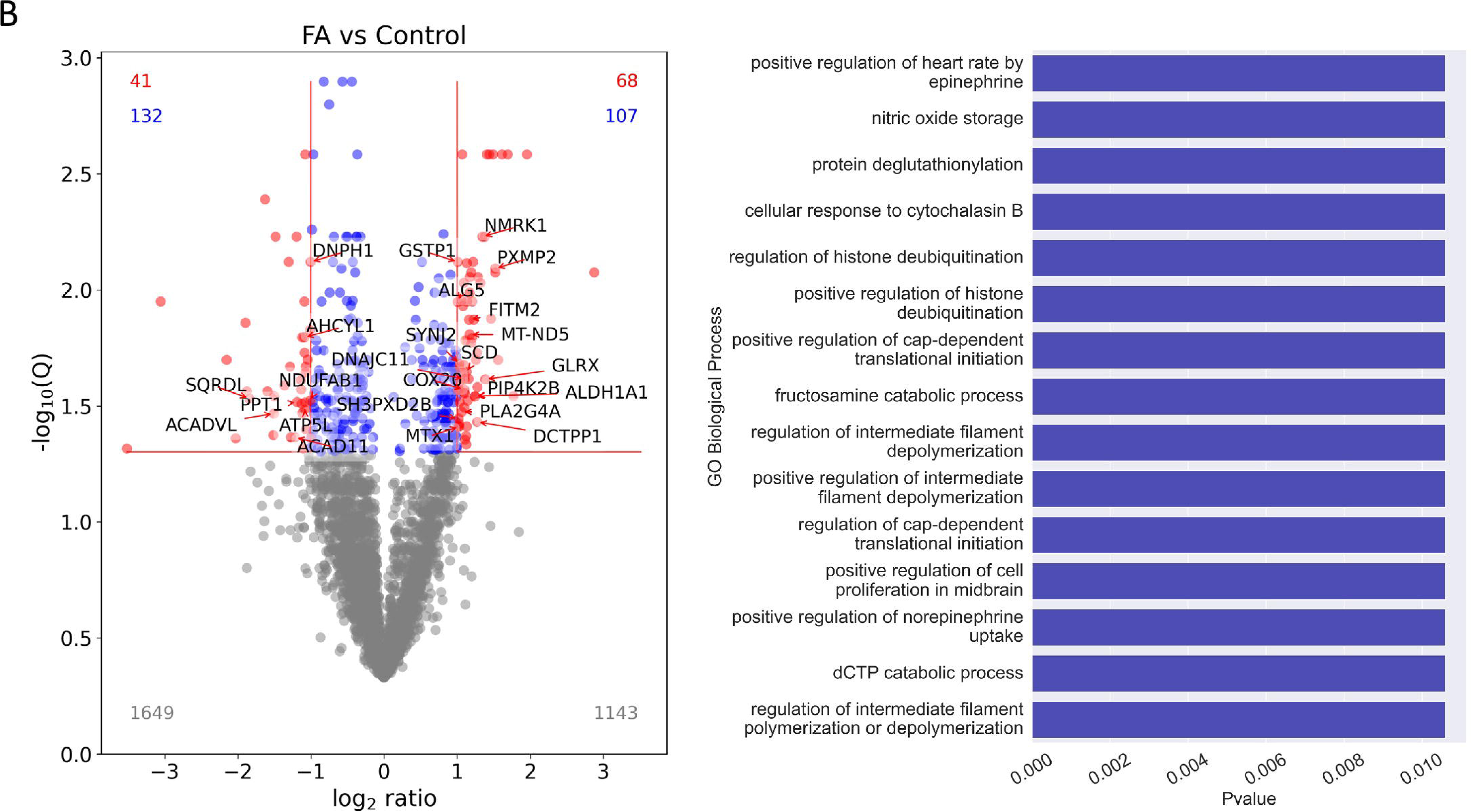
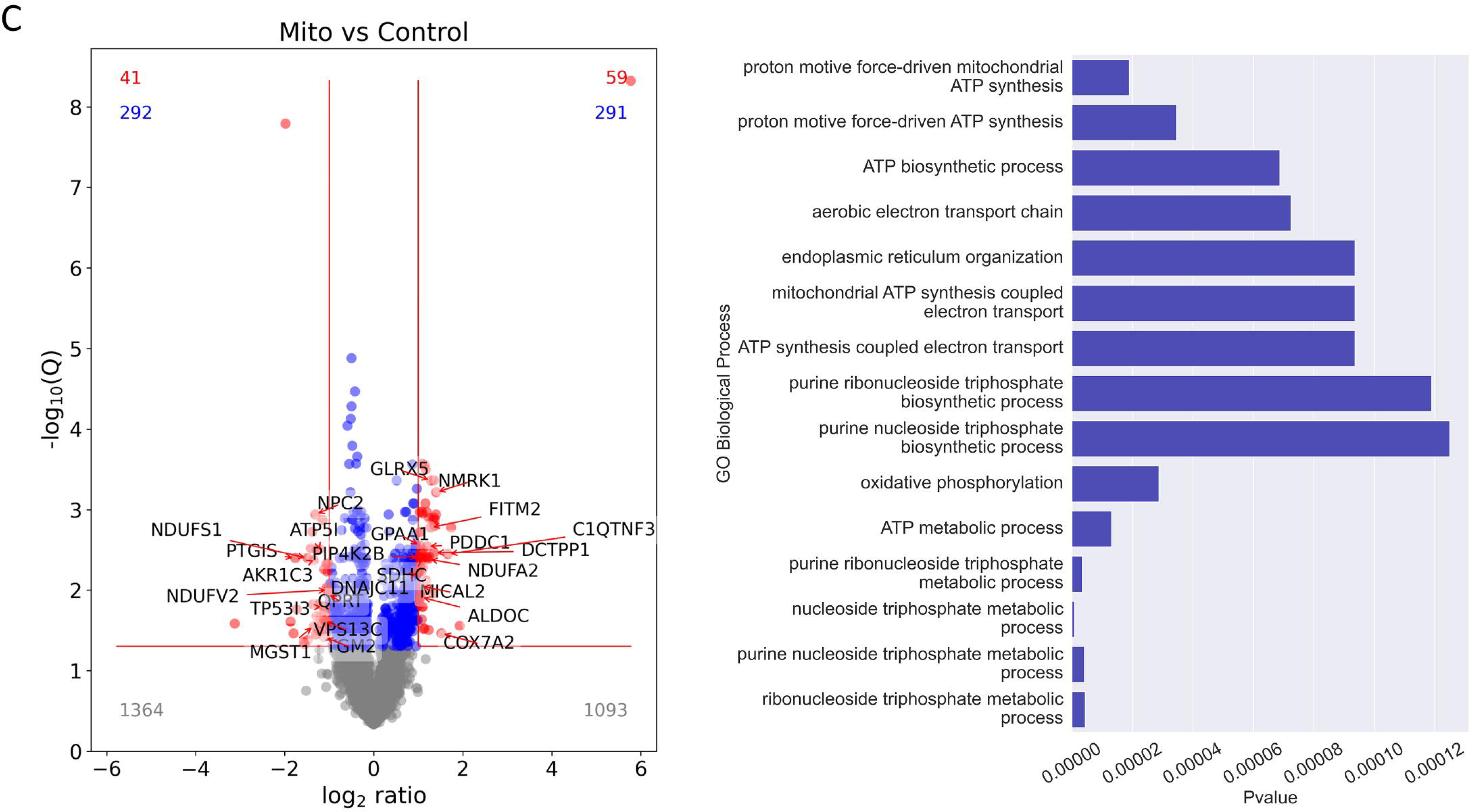
Proteomics identification of differentially expressed (regulated) proteins and corresponding functional over-representation analysis of enriched GO terms for (A) OA, (B) FA and (C) Mito IEM disease patient groups. Volcano plots demonstrate the distribution of log2 fold change and FDR-corrected statistical scores for all quantified proteins. Red dots correspond to regulated proteins that are further used for the GO analysis. The number of proteins in the respective areas of the volcano plot are mentioned using the corresponding colors. Regulated proteins with experimentally annotated functions in metabolic processed are highlighted. With respect to the GO analysis, the GO BP ‘Slim’ dataset was used for generating the results shown for OA, and the GO BP ‘Complete’ dataset was used for the FA and Mito patient groups. Enriched GO BP terms were identified, sorted according to decreasing Fold Change values; the Top 15 terms were then plotted.

To address these functional implications more closely, we performed a statistical over-representation analysis based on Gene Ontology (GO) Biological Process (BP) terms, as implemented by PANTHER^15^. GO BP functional terms associated with the significantly regulated proteins in each disease category were compared against annotations for the complete human genome. Top over-represented terms were sorted based on their fold enrichment and the statistical scores of the top 15 enriched terms are shown in **Figure 1**. Several of these identified GO BP terms closely mirror known patient clinical features as well as expected perturbed pathways, such as ATP synthesis and electron transport oxidative phosphorylation for Mito patients and amino acid biosynthetic processes for OA patients.

Unlike the differentially expressed (DE) proteins and the subsequent GO analysis which provide several initial clues to derive hypothesis behind the IEM etiology, a PCA performed on the data however, showed a relatively large overlap between the disease groups (and even with the control group), indicating that the overall protein expression profiles were relatively similar (**Figure S1**). The high inter-sample similarity between the global protein expression patterns were also demonstrated by a correlation matrix (**Figure S2**). This was not very surprising given the fact that the impairment of mitochondrial function has been implied in all of their pathophysiologies. Moreover, in general, only one protein is affected or defective in IEM, whether that’s based on bi-allelic genetic variants or dominant mono-allelic, mtDNA or hemizygosity. Nevertheless, as the identity of the DE proteins and the lists of over-represented biological functions showed, interesting patterns of differences began to emerge. Moreover, as we describe later, even a single SNP can and does translate into significant differences at the level of the cellular assemblies that help pinpoint perturbed biological functions and the underlying causative molecular machinery.

### Transcriptomics data exhibits low overall correlation with proteomics

Similarly, the transcriptomics data was processed to explore disease specific differences. Following processing and filtering steps (see Methods), we quantified 16202 transcripts across the disease groups and identified 782, 190 and 427 among those to be differentially regulated for the OA, FA and Mito groups, respectively, relative to control samples, based on FDR <= 0.05 and a fold change cutoff of 2. Next, we calculated the correlation between the log2FC values at the protein and transcript levels: the Pearson correlation coefficients were 0.33, 0.13 and 0.21 for the OA, FA and Mito categories, respectively, indicating low quantitative matches between the measurements at these two omics levels. Several recent large-scale studies also demonstrated the low overall concordance between RNA and protein level information from mouse data ^2,3^ as well as in human tissues leading to the necessity of obtaining protein level information to better explain genetic disease phenotypes ^1,4^.

We also performed a GO BP term over-representation analysis, using the differentially expressed gene lists, as done for the proteomics data. Given that the number of differentially expressed genes were larger in comparison to the numbers of differentially expressed proteins, we obtained greater statistical power in the GO BP analysis, in terms of the FDR values observed, for example. As a result of the larger gene sets, we also observed a greater number (and functional variety) of over-represented terms; briefly summarized, the most relevant ones included metabolic functions including several generalized lipid metabolic regulation terms (for OA, FA and Mito categories), prostaglandin and prostanoid biosynthesis, unsaturated fatty acid biosynthesis and cyclooxygenase pathway terms (in the OA and Mito groups).

When a PCA was used to visualize overall trends in the data (**Figure S3**), mirroring the observations from the proteomics level, here also we found the disease groups to be very similar overall indicating that both at the transcript and protein levels inter-sample expression profiles were relatively similar. One potential explanation is that, given that all the samples were fibroblasts, this particular cell type may have limited options for differentiation, irrespective of the underlying genetic alterations. Additionally, the culturing conditions employed (refer to Methods) were formulated to maintain the metabolic stability of fibroblasts, whereas individuals with inborn errors of metabolism (IEM) typically experience symptoms when faced with metabolic challenges.

### Analysis of untargeted metabolomics data reveals several lipids participating in glycerophospholipid metabolism as the top discriminatory features

The LC-MS based untargeted metabolomics dataset provided a total of 5644 features post-processing (see Methods). Similar to the omics datasets described above, a PCA performed on all detected compound ions showed large overlaps between the IEM groups (**Figure S4**). Next, we identified the top 200 significant features based on an ANOVA statistical test (p<0.05) and used MS-FINDER to annotate the corresponding compounds (see Methods for details). Based on the ANOVA scores and a visual inspection of the data, we found that several lipid species were significantly different across the cohort. One of the most striking differences were observed for the two related classes of 2-acyl-sn-glycero-3-phosphoethanolamines and 1-acyl-sn-glycero-3-phosphoethanolamines, which showed a large increase in FA and Mito samples, and particularly in the Mito group (compared to both Control and OA samples) (**Figure 2**). Both these classes of molecules are linked to the biosynthesis of phosphatidylethanolamine (PE), a major inner membrane phospholipid. PE is linked to several important metabolic processes including energy metabolism, cell signaling cascades as well as providing the phosphoethanolamine moiety of glycosylphosphatidylinositol (GPI) anchors, necessary for membrane anchoring of several proteins^16^. Within the context of glycerophospholipid metabolism, 1-acyl-sn-glycero-3-phosphocholines also showed the same trend, i.e., higher levels were found in FA and Mito groups, and particularly in the latter (**Figure 2**). These are metabolic derivatives of phosphatidylcholines and thereby also linked to PEs. Moreover, 1-acyl-sn-glycero-3-phosphocholines are also connected to sn-glycerol-3-phosphate (G3P), a metabolite which branches off from the main glycolytic pathway through the conversion of dihydroxyacetone phosphate (DHAP) by the enzyme G3P dehydrogenase and provides the backbone onto which the fatty acids are connected. In a previous study, it was also shown that mitochondrial related IEM showed higher use of the glycerol 3 phosphate shuttle^17^.

**Figure 2.**
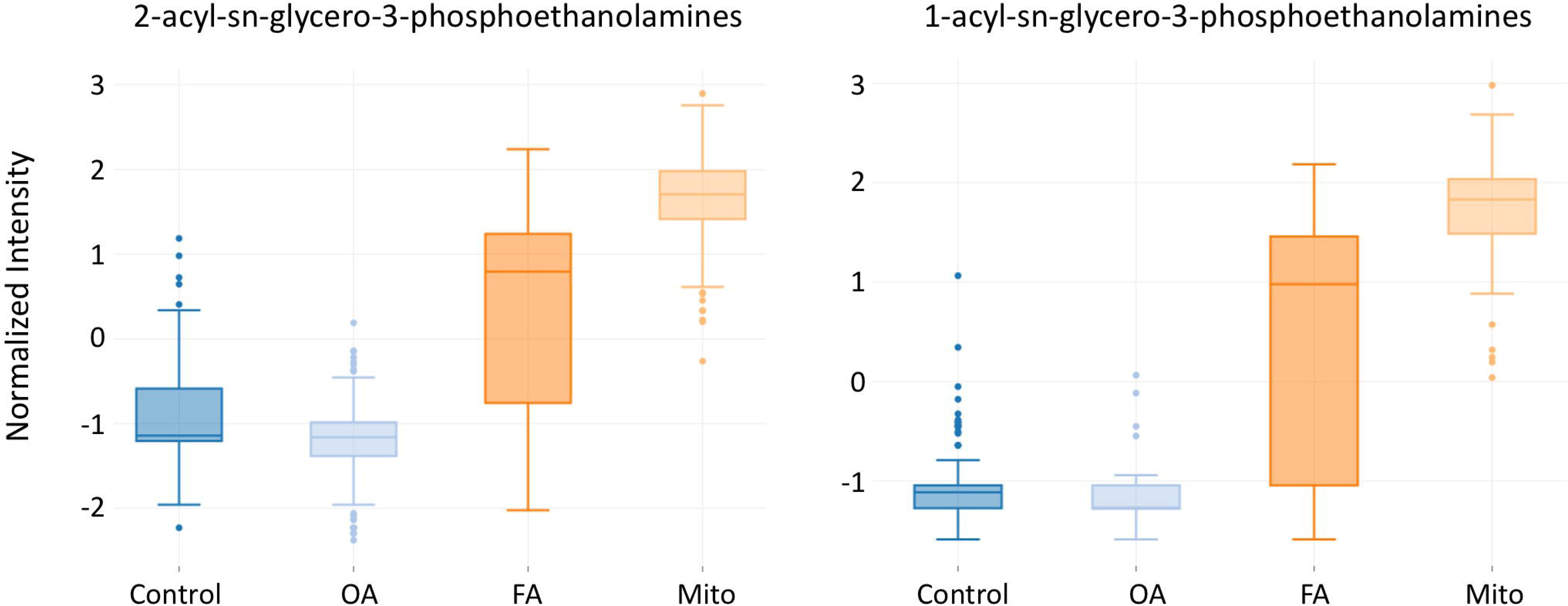
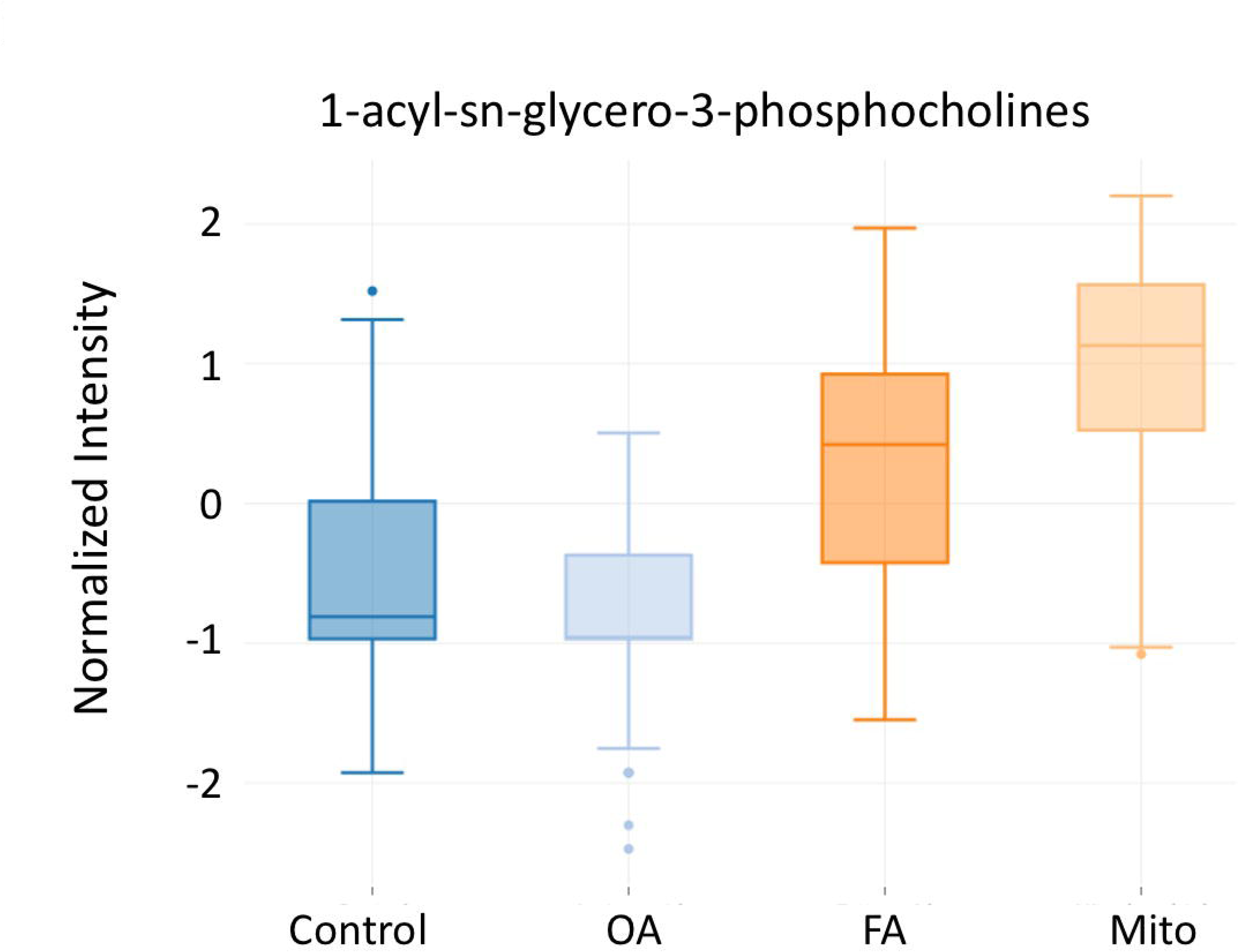
Relative abundances (shown as normalized intensities on the y-axis, measured using untargeted metabolomics) of selected lipid species that showed the largest changes between the IEM disease groups. The annotated lipid molecules in each of the three compound categories were as follows: 2-acyl-sn-glycero-3-phosphoethanolamines (LysoPE(0:0/24:6(6Z,9Z,12Z,15Z,18Z,21Z)), LysoPE(0:0/20:4(5Z,8Z,11Z,14Z)), LysoPE(0:0/22:4(7Z,10Z,13Z,16Z)) and LysoPE(0:0/22:5(4Z,7Z,10Z,13Z,16Z))); 1-acyl-sn-glycero-3-phosphoethanolamines (LysoPE(22:5(4Z,7Z,10Z,13Z,16Z)/0:0), LysoPE(22:6(4Z,7Z,10Z,13Z,16Z,19Z)/0:0) and LysoPE(22:5(7Z,10Z,13Z,16Z,19Z)/0:0)), and, 1-acyl-sn-glycero-3-phosphocholines (LysoPC(20:3(5Z,8Z,11Z)), LysoPC(20:4(5Z,8Z,11Z,14Z)), LysoPC(22:5(4Z,7Z,10Z,13Z,16Z)) and LysoPC(22:6(4Z,7Z,10Z,13Z,16Z,19Z))).

Interestingly, several studies have demonstrated the transport of phospholipids (PS, PE and PC) between the ER and mitochondrial membranes at locations where the ER and outer mitochondrial membranes are closely apposed physically. In yeast, the ER-mitochondria encounter structure (ERMES) complex is responsible for tethering these two organelles and mutations in subunits of the ERMES complexes disrupted phospholipid biosynthesis^18^. In metazoans, orthologues of the ERMES complex have not been found, however, the *VPS13* gene family that encodes lipid transfer proteins, are known to be highly conserved from yeast to higher eukaryotes and can possibly compensate for the absence of ERMES. Looking into the proteomics dataset, we found that VPS13C was significantly downregulated in the Mito samples (log2FC -1.12, FDR 0.02; **Figure 1C**), but not in any of the other groups. The other VPS13 paralogs were not quantified in the proteomics dataset. It is interesting that VPS13C abundance was significantly different only in Mito samples, which also showed the largest relative increases in the phospholipids mentioned above. VPS13C tethers the ER to the mitochondrial outer membrane^19^, although other studies also suggested that VPS13C has a major role in ER-late endosome/lysosome contacts and lipid transfer^20^. VPS13C depletion affected mitochondrial morphology and renders mitochondria more vulnerable to stress^19^.

Altogether, the multi-omics data point towards a close biochemical relationship between the FA disorders, mitochondrial disorders, and lipids. Biochemically, both these classes of IEMs are indeed closely connected as the oxidation of FAs occurs within the mitochondria and provides acetyl-CoA, a key intermediate of the Krebs cycle generating the essential cofactors (electron donors) for the oxidative phosphorylation chain. In this respect, lipids are also closely connected to mitochondrial IEM as lipid hydrolysis provides FA which in turn can be oxidized towards acetyl-CoA.

### Dissecting perturbations in specific protein complexes elucidates disease mechanisms

In order to pinpoint exactly the molecular machinery that is perturbed in IEM patients, we applied an approach based on the analysis of the human ‘complexome’ (i.e., the complete complement of experimentally validated protein assemblies that are currently known in this species). Specifically, we queried proteome-wide, manually curated datasets of human protein complexes/assemblies from the Complex Portal resource^12^ and identified those complexes in which one or more subunits were found to be significantly regulated based on the proteomics data.

We employed two approaches in parallel: firstly, using a local network approach and secondly, a global approach based on interrogating the entire human complexome. In the first approach, we used the known mutated gene information for every patient (**Table 1**) to extract the direct protein-protein interaction (PPI) partners of each affected protein from the global human interactome; this local PPI network was then queried for protein complex memberships (a schematic representation is shown in **Figure 3A**). This local network analysis provides an idea of the immediate functional and physical interaction environment of the mutated protein and can highlight complexes that are highly relevant to the disease under investigation. For example, several subunits of mitochondrial respiratory chain complexes were identified as being perturbed in 5 out of 6 Mito and 1 out of 4 FA patients (**Table 2**). One limitation of this approach is that the analysis is dependent on the number of experimentally known PPI partners of the mutant protein of interest. If these are relatively few (based on information currently available in PPI databases), it limits the usefulness of the local network-based approach. In this cohort, the average number of experimentally known PPI partners were 9.5, 25 and 43.7 for the mutated proteins of the OA, FA and Mito patient groups, respectively (see also **Table 4**).

**Figure 3.**
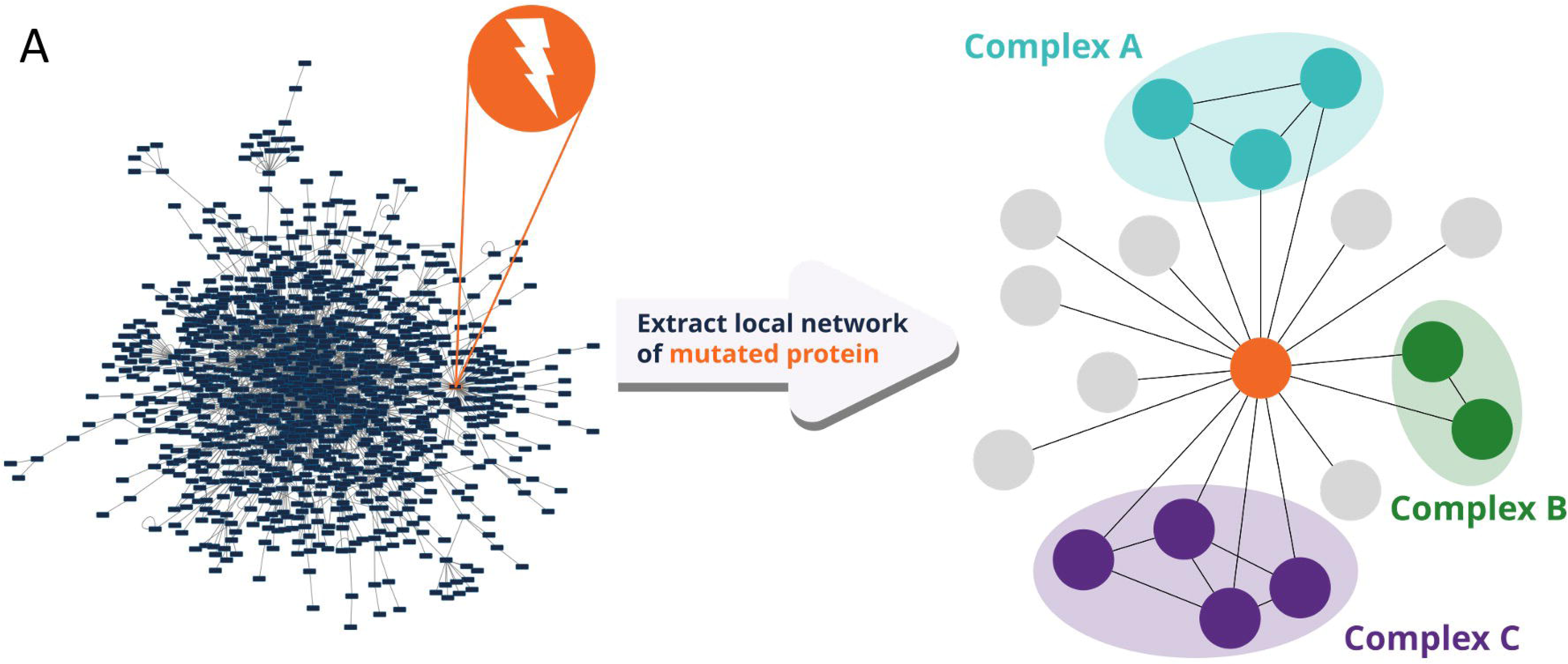
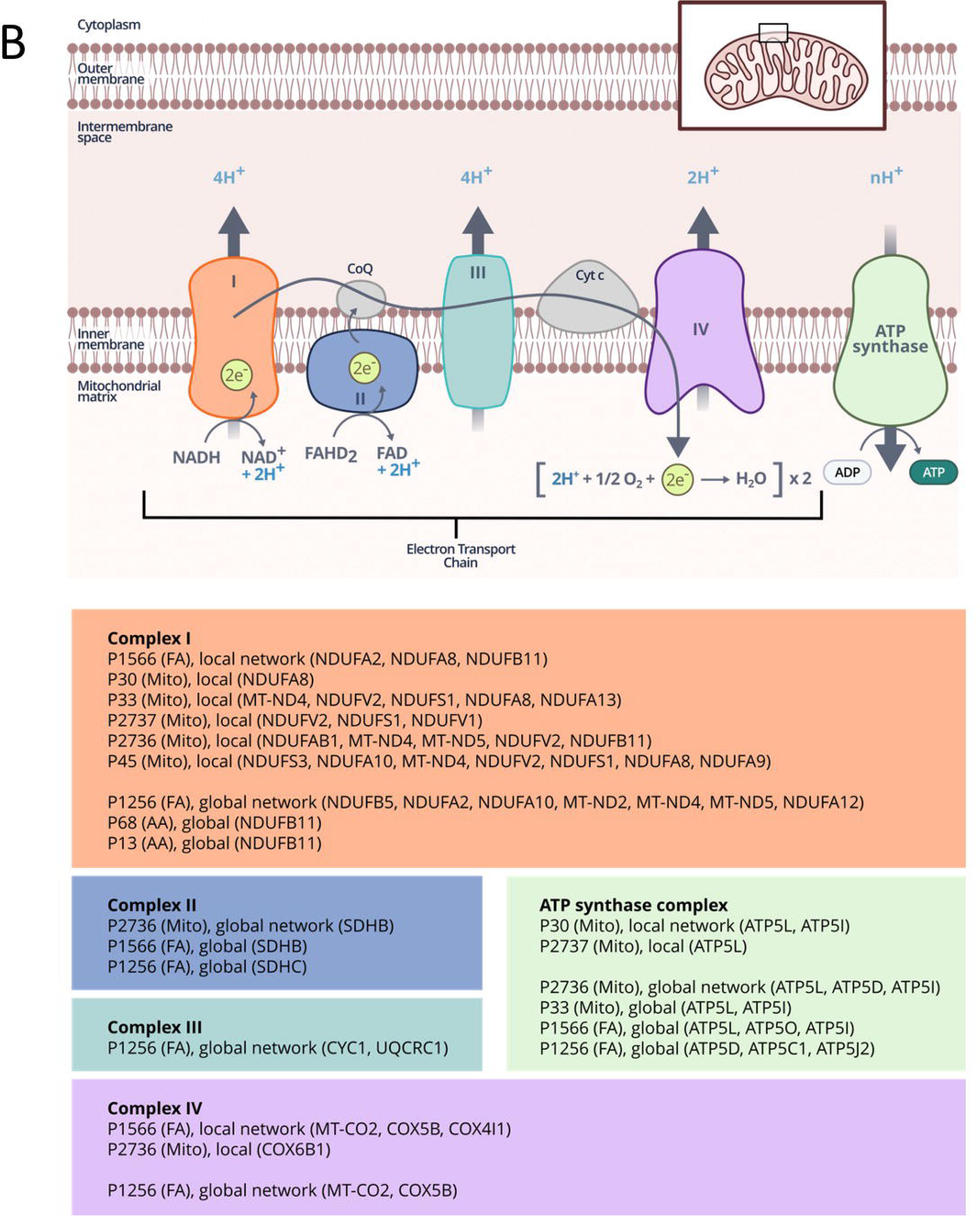
Local network is identified for each known mutated gene, by zooming in from the global PPI network, as shown in (A). Having extracted the local network of the mutated protein, memberships of protein complexes are then queried. (B) Schematic diagram of the mitochondrial electron transport chain (ETC). Across the sample cohort studied here, mitochondrial ETC complexes were consistently found to be perturbed in IEM samples vs controls. For each complex (color-coded) we detail the patient samples in which the respective ETC complexes were affected, the identity of the subunits that were perturbed and whether the identification was made from the local network or the global complexome analysis (shown on the figure as ‘local’ or ‘global’, respectively).

**Table 2.**
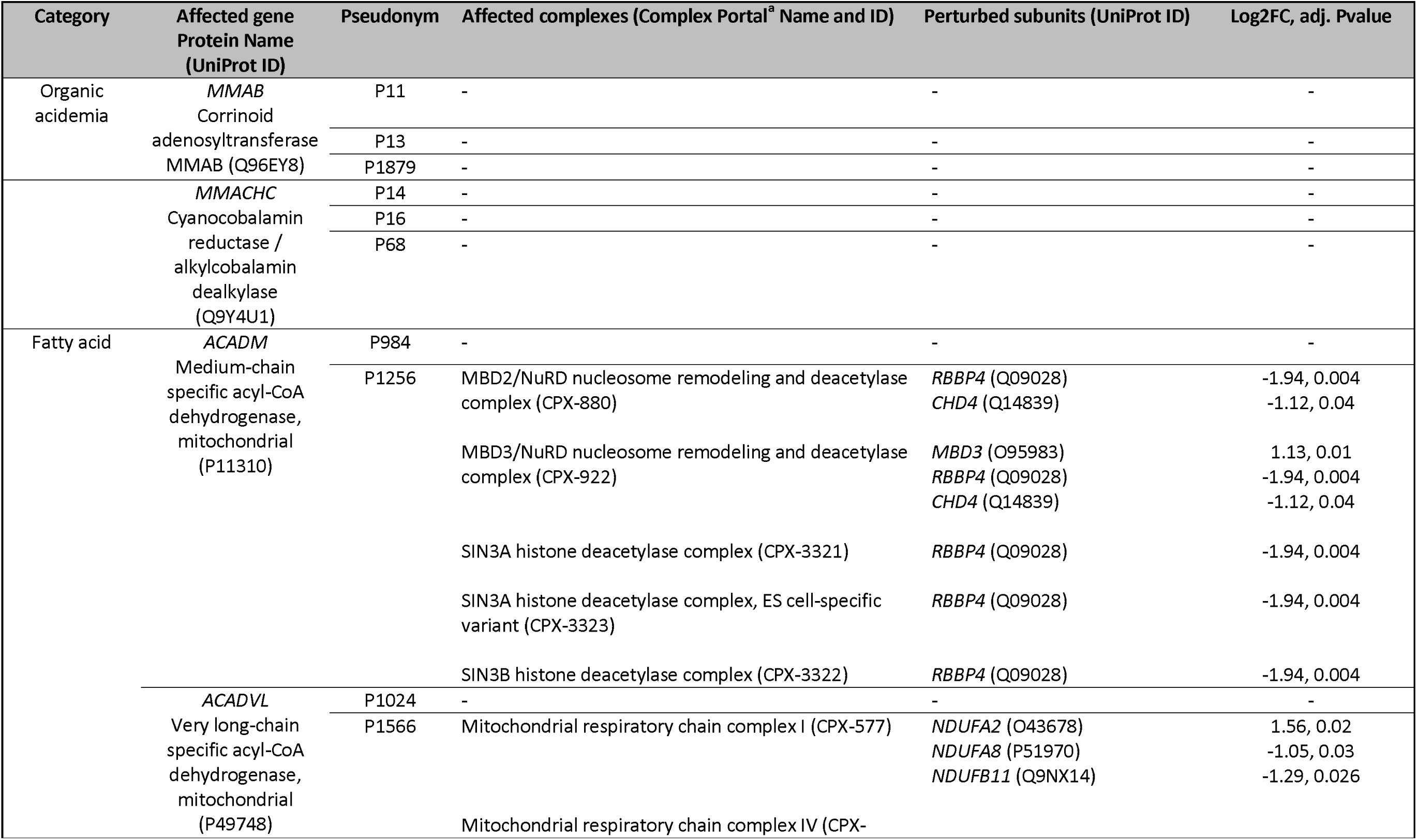
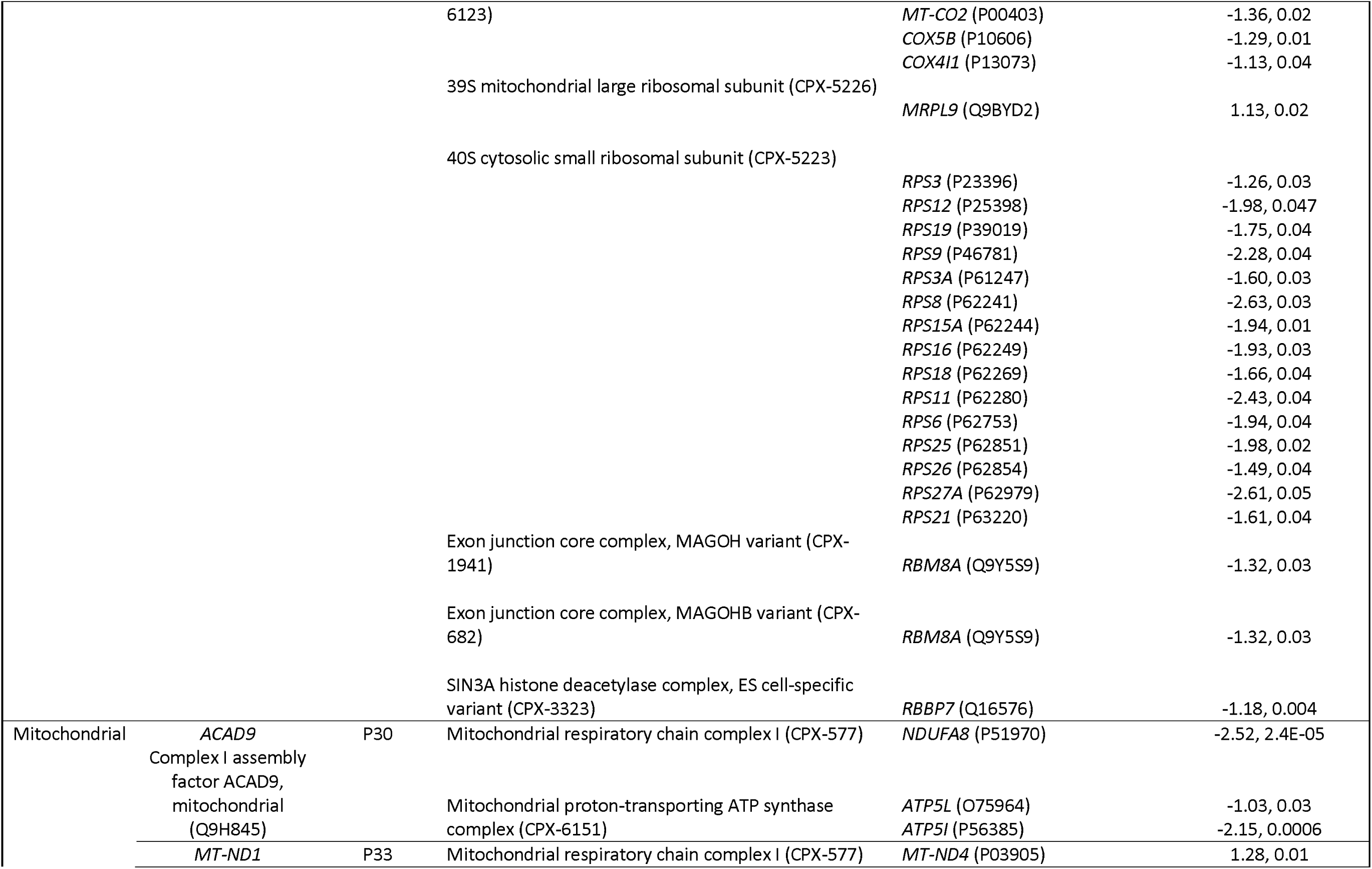
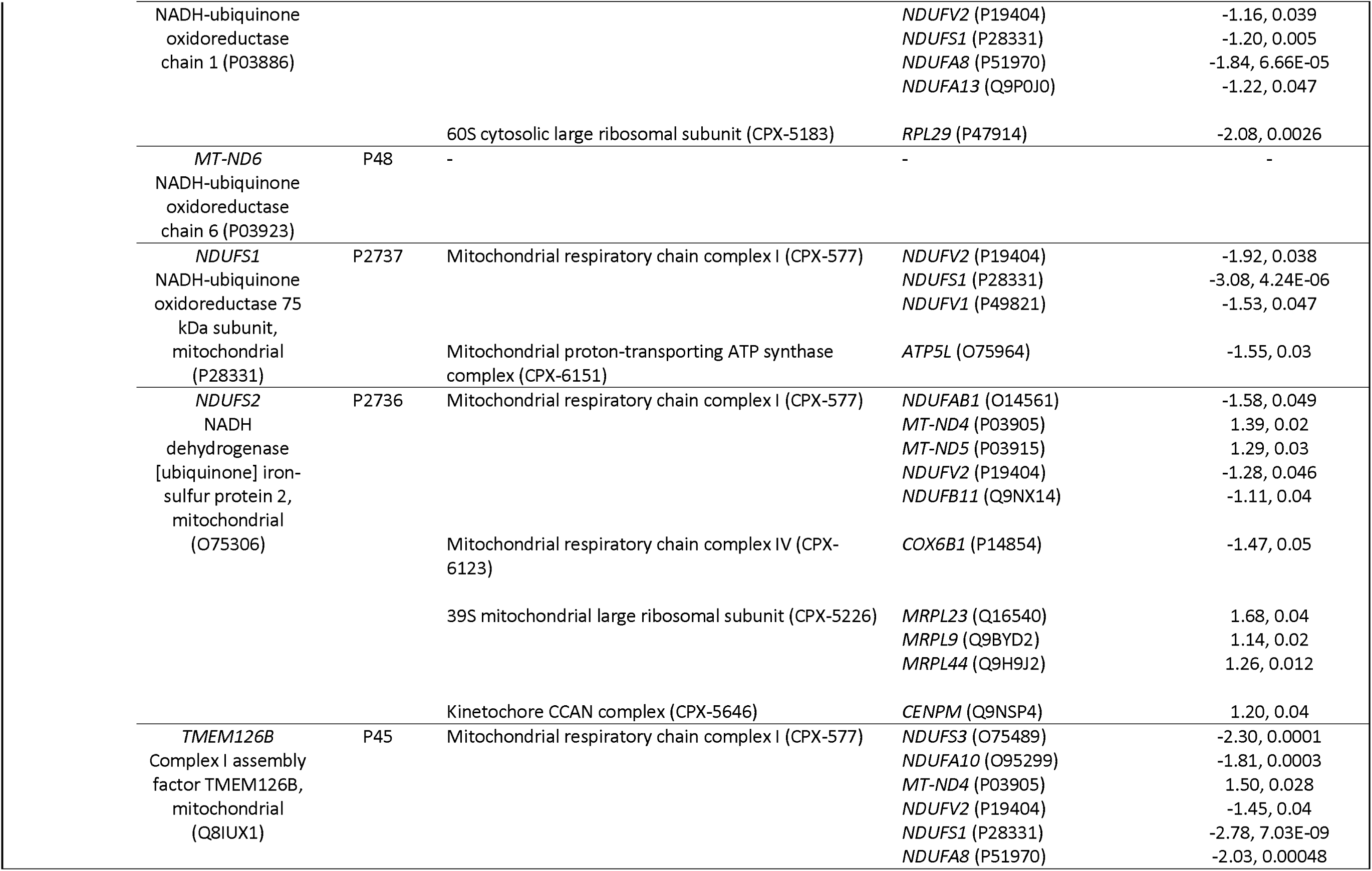
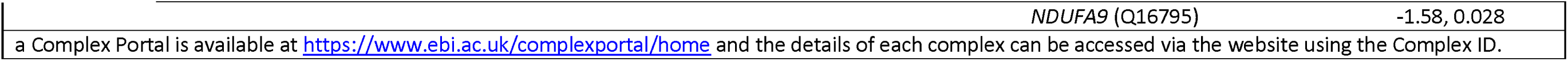
Summary of affected protein complexes per patient sample (based on the proteomics measurements overlaid onto the local PPI network of each mutated gene).

**Table 3.** Complexes with histone (de)acetylase and chromatin remodeling functionality for samples showing upregulation of histone acetylation levels compared to control groups.

| **Complex Name**<br>**(Complex Portal ID)** | **Affected in patient**<br>**ID(s)** | **Perturbed subunit(s) within the complex (Log2FC,**<br>**FDR)** |
| --- | --- | --- |
| WICH chromatin remodelling complex<br>(CPX-757) | P30<br>P45<br>P68 | BAZ1B (Q9UIG0) (1.43, 0.05)<br>BAZ1B (Q9UIG0) (2.05, 0.02)<br>SMARCA5 (O60264) (-2.56, <0.0001) |
| B-WICH chromatin remodelling complex<br>(CPX-1099) | P30<br>P45<br>P68 | BAZ1B (Q9UIG0) (1.43, 0.05)<br>BAZ1B (Q9UIG0) (2.05, 0.02)<br>SMARCA5 (O60264) (-2.56, <0.0001) |
| MBD2/NuRD nucleosome remodeling and deacetylase<br>complex<br>(CPX-880) | P45 | CHD4 (Q14839) (-1.41, 0.04) |
| MBD3/NuRD nucleosome remodeling and deacetylase<br>complex<br>(CPX-922) | P45 | CHD4 (Q14839) (-1.41, 0.04) |
| NoRC chromatin remodelling complex<br>(CPX-432) | P68 | SMARCA5 (O60264) (-2.56, <0.0001) |
| ACF chromatin remodeling complex<br>(CPX-434) | P68 | SMARCA5 (O60264) (-2.56, <0.0001) |
| RSF complex<br>(CPX-455) | P68 | SMARCA5 (O60264) (-2.56, <0.0001) |
| CHRAC chromatin remodeling complex<br>(CPX-785) | P68 | SMARCA5 (O60264) (-2.56, <0.0001) |
| NSL histone acetyltransferase complex<br>(CPX-809) | P45 | HCFC1 (P51610) (-3.62, 0.03) |

**Table 4.** Pearson’s correlation coefficients (PCC) calculated between the proteomics and transcriptomics datasets per sample globally and for subsets of protein-gene pairs therein.

| Sample | Global PCC | PPI partners of mutated protein <sup>a,e</sup> | Local network complexes <sup>c,e</sup> | Global network complexes <sup>d,e</sup> |
| --- | --- | --- | --- | --- |
| P11 | 0.31 | -0.13 (6/17) | -0.12 (15/56) | 0.09 (672/2226) |
| P13 | 0.29 | 0.91 (6/17) | -0.04 (15/56) | 0.20 |
| P1879 | 0.18 | 0.78 (6/17) | 0.33 (15/56) | 0.06 |
| P14 | 0.13 | - (2/2) <sup>b</sup> | - | 0.02 |
| P16 | 0.16 | - (2/2) <sup>b</sup> | - | 0.05 |
| P68 | 0.24 | - (2/2) <sup>b</sup> | - | 0.13 |
| P984 | 0.21 | 0.55 (4/14) | 0.44 (10/40) | 0.04 |
| P1256 | 0.06 | -0.45 (4/14) | 0.21 (10/40) | 0.11 |
| P1024 | 0.09 | 0.16 (12/36) | 0.17 (93/216) | 0.17 |
| P1566 | 0.06 | -0.24 (12/36) | 0.14 (93/216) | 0.07 |
| P30 | 0.19 | 0.36 (28/79) | 0.04 (69/168) | 0.16 |
| P33 | 0.18 | -0.04 (12/30) | 0.006 (67/108) | 0.09 |
| P48 | 0.06 | 0.99 (3/5) | 0.04 (28/55) | 0.08 |
| P2737 | 0.20 | 0.03 (29/79) | 0.098 (77/201) | 0.067 |
| P2736 | 0.09 | 0.004 (26/58) | -0.16 (61/167) | 0.01 |
| P45 | -0.002 | -0.65 (4/11) | -0.18 (28/55) | -0.08 |

a Mutated protein per sample is shown in Table 1.
b Correlation coefficient not shown because only 2 data points.
c Local network complexes are detailed in Table 2.
d The complete human complexome from Complex Portal
(https://www.ebi.ac.uk/complexportal/home).
e Numbers shown in parentheses: the denominator indicates the number of proteins within that category, while the numerator is the number of proteins from that category for which both proteomics and corresponding transcriptomics data were quantified.

Therefore, to obviate this issue and, more importantly, to obtain a systems-wide overview of possible perturbations triggered by IEM, the proteomics data were mapped onto the entire complexome. For *Homo Sapiens*, the October 2021 release (used here) of the Complex Portal contained 1256 multiprotein assemblies. Complexes with at least one perturbed subunit (defined as proteins with a log2FC > 1 or <-1 and FDR<=0.05) were identified (**Tables S1**; presented as .csv files) per patient sample. All the perturbed complexes identified earlier from the local network analysis were also recapitulated here. On average, the number of perturbed complexes were 7, 121 and 56 for the OA, FA and Mito categories, respectively. Manual inspection of individual complexes per sample highlighted novel and potentially critical, disease-relevant perturbed complexes. For instance, all the complexes forming the mitochondrial electron transport chain (ETC) were consistently affected in multiple samples from the Mito and FA patient cohorts (**Figure 3B**) which might explain the poor separation on the PCA plots. Oxidative phosphorylation (OXPHOS) is a key cellular pathway localized in mitochondria that is responsible for cellular energy generation and it comes as no surprise that this is massively affected in IEMs^21^, particularly in the FA and Mito samples, as shown by the proteomics data.

OXPHOS is a complex pathway that includes the redox reactions mediated by the ETC complexes that generate the proton motive force ultimately used for ATP synthesis. Moreover, the correct function of OXPHOS also requires the orchestrated synthesis of the numerous subunits (that are encoded both by the nuclear as well as the mitochondrial DNA), mitochondrial import of the nuclear encoded subunits, their integration into the inner mitochondrial membrane, and finally, their coordinated assembly pathways which also involves the formation of a number of characterized assembly intermediates. Based on our global complexome analysis (details provided in **Tables S1**), we indeed saw many of these associated, wider mitochondrial functions affected in multiple Mito and FA samples. For example, other perturbed mitochondrial assemblies included the TIM22 mitochondrial inner membrane twin-pore carrier translocase complex (CPX-6124, in sample P1256), the TIM23 mitochondrial inner membrane pre-sequence translocase complexes (CPX-6129 and 6130, in sample P45), mitochondrial processing peptidase complex (CPX-6243, in P33 and P1566), the SAM mitochondrial sorting and assembly machinery complex (CPX-6133, in P2736, P1256 and P1566), the TOM40 mitochondrial outer membrane translocase complex (CPX-6121, in P1256 and P1566), several variants of the mitochondrial intermembrane space protein transporter complexes (CPX-6125, -6126, -6131 and -6132; in sample P1256), the MICOS mitochondrial contact site and cristae organizing system complex (CPX-6141, in P1256), mitochondrial NIAUFX iron-sulfur cluster assembly complex (CPX-5641, in P2736), mitochondrial BOLA1-GLRX5 and BOLA3-GLRX5 iron-sulfur cluster assembly complexes (CPX-6862 and CPX-6863, in P1256 and P1566), and, several subunits of the 28S and 39S mitochondrial ribosomal subunits (in P33, P45, P2736, P1256 and P1566). We also remark that significant changes in phospholipid content and composition, such as of PE, may also lead to abnormal mitochondrial membrane architecture and therefore the complexes they contain also are affected.

Further, to summarize the information, we computed the top 20 most frequent GO BP terms linked to the perturbed complexes identified from the global complexome analysis (**Figure S5**). Interestingly, one of the functional categories that appeared to be consistently perturbed, particularly in the FA and Mito groups, were complexes involved in histone (de)acetylation and associated chromatin remodeling functions. These complexes included major HDAC complex families including NuRD and SIN3 and chromatin remodeling WICH and SWI/SNF complexes (**Tables S1**). Since these complexes play major roles in myriad facets of transcription regulation, we decided to explore this aspect in more detail. To that end, we performed an experimental assay to determine changes in the acetylation state of histones in those specific samples with significant numbers of perturbed complexes having this functionality.

### Histone acetylation is affected in FA and Mito IEM

Using a pan-acetylation antibody on purified histones from the healthy and IEM fibroblasts we investigated the hypothesis derived from the complexome observations that histone acetylation would be affected in certain samples. **Figure 4** shows the results from the Western Blot which indeed validated the prediction that histone acetylation would be perturbed in those patient cell lines. Especially within the Mito group, P0030 and P0045 showed increased levels of histone acetylation while P0033 and P2736 displayed reduced acetylation levels. In order to rationalize these observations based on the global complexome perturbation analysis (**Tables S1**), we looked at the specific affected complexes with relevant histone (de)acetylation functionality in the samples P30, P45 and P68 (**Table 3**) which showed elevated histone acetylation levels (**Figure 4**).

**Figure 4.**
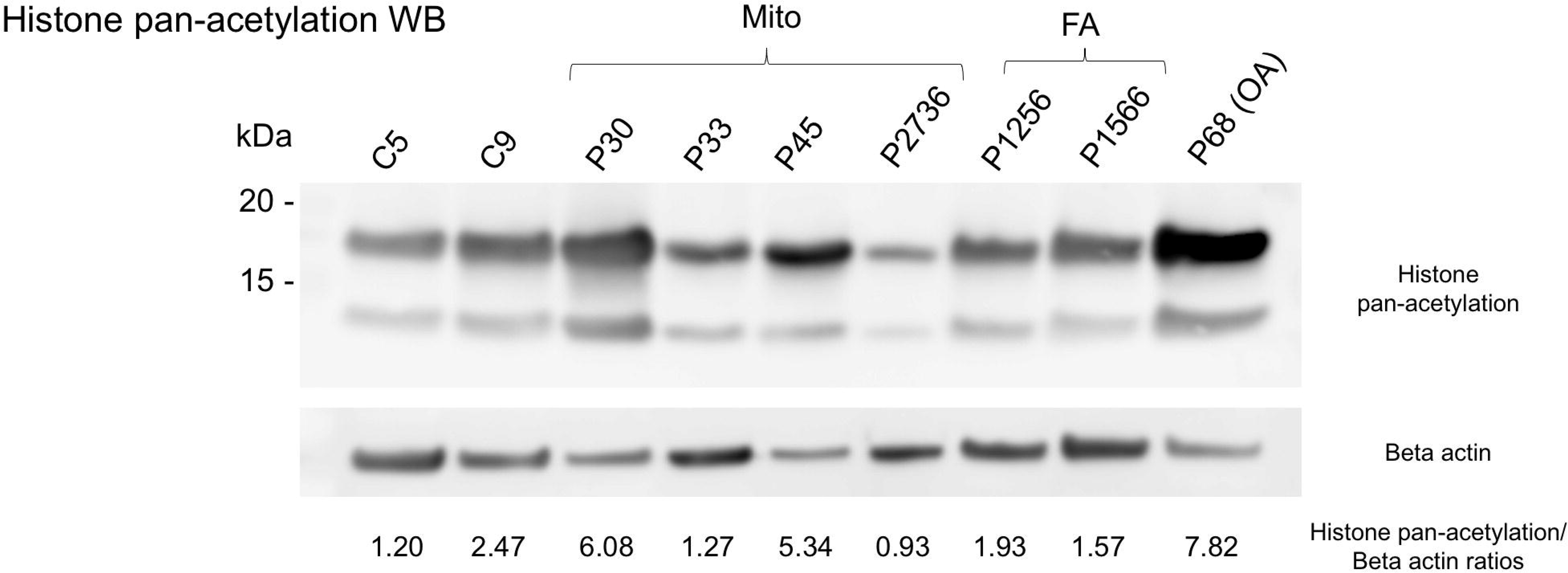
Histone acetylation assays for selected samples (controls and IEM samples as marked on the plot). Increased levels of histone pan-acetylation in several patient fibroblasts were identified. Isolated histone extracts were separated on a 10% Bis-Tris gel, transferred to a nitrocellulose membrane, and blotted with the corresponding antibodies. Beta actin was used as a loading control. Densitometric quantifications are shown below each lane in the form of the ratio of the combined histone pan-acetylation signals divided by the signal from beta actin.

In both P30 and P45 samples, the BAZ1B nuclear protein, with a key role in chromatin remodeling through its participation in two distinct complexes (WICH and B-WICH), is upregulated. The WICH complex localizes to DNA replication sites and increases chromatin accessibility for replication and epigenetic factors including chromatin modifying enzymes^22^. The B-WICH complex facilitates the recruitment of histone acetyltransferases (HATs) to rDNA promoters and is involved in H3K9 acetylation^23^. In fact, BAZ1B knock down causes both lowered HAT recruitment to promoters and global histone acetylation, specifically H3K9-Ac, to be decreased^23^. P45 also has lowered levels of the CHD4 ATPase subunit of MBD2 and MBD3/NuRD complexes (**Table 3**). Both these complexes function by removing acetyl group from nuclesosomes and therefore reduced acitivity is in accordance with higher histone acetylation levels (**Figure 4**). Taken together, these observations potentially explain mechanistically how higher BAZ1B (in P30 and P45) and lower CHD4 (in P45) levels may translate into a concomitant increase in global histone acetylation levels in both samples. Also interestingly, in the P45 sample, but not in P30, we saw a sharp downregulation of the HCFC1 protein which forms part of a number of histone lysine N-methyltransferase (HMT) complexes (**Table S1**). In addition, HCFC1 also associates with the Sin3 histone deacetylase (HDAC)^24^ and their decreased association because of decrease in HCFC1 levels could form an additional, synergistic effect and explain the large increase in histone acetylation that we observed for P45 (**Figure 4**).

In the P68 sample, we found that the SMARCA5/SNF2H ATPase motor protein was significantly downregulated (**Table 3**). SMARCA5 forms the catalytic core around which multiple accessory subunits (such as BAB1B) can attach in a cell type-dependent, combinatorial fashion, resulting in the formation of multiple, polymorphic chromatin remodeling complexes^25^, all of which are potentially affected by the perturbation of the central ATPase motor subunit (**Table 3**). For example, the NoRC chromatin remodelling complex (CPX-432) recruits HDACs and HMTs, leading to heterochromatin formation and the repression of ribosomal gene transcription. SMARCA5 downregulation and therefore lowered NoRC functions, particularly the HDAC functionality, is in line with the higher acetylation observed in P68 (**Figure 4**).

The histone code is intricate and complex combinatorics are employed in the regulation^26^ and this is reflected in the number of different histone modifying HAT, HDAC and HMT complexes (and their variants) that are currently known and that were identified in our global complexome analysis as being perturbed (see the csv output files for P30, P33, P45, P2736, P1256, P1566 and P68; **Tables S1**). Despite the complexity, nevertheless we can harness the proteomics measurements and the associated complexomics analysis to derive insights into possible mechanistic explanations into changes in histone acetylation status. Although such discovery studies can be taken much further, here our motivation was to demonstrate how starting from the genetic mutation, looking at the results going upto the level of dysregulated protein complexes can be a valuable tool for multi-omics studies as a discovery tool predicting functional consequences and generating new hypotheses that can be further experimentally validated.

### Dissecting protein-transcript abundance change correlations at the level of the interaction neighborhood of the mutated genes and associated complexes

Understanding the relationship between protein and mRNA levels is challenging given the dynamic changes required for cellular adaptation under diverse stimuli as well as resulting from genotypic variation. This complexity arises from the multitude of regulatory processes that independently govern the production, localization, activity, availability and degradation of the transcriptome and the proteome, leading to the discrepancy commonly observed between mRNA and protein levels^27^. Here also, we first asked to what extent the observed changes in protein abundances correlated with the corresponding changes in transcript-level abundances. Overall, this was poor to moderate for all samples studied (**Table 4**). This is a common problem in multi-omics as has also been demonstrated in several recent large-scale studies^1,2,4^. However, in order to parse out whether certain functional molecular subsets might be more synergistically regulated, we computed mRNA-protein abundance change correlations for the direct PPI partners of each mutant protein as well as for the subunits of complexes that were identified as being a part of the local PPI network for that protein. Experimentally validated PPI partners were obtained from the IntAct database^11^ and local network complexes were identified as shown earlier in **Figure 3A**. When considering direct PPI partners, we observed in several cases an improved correlation between mRNA-protein abundance changes (compared to the global correlation), going up to 0.9 (**Table 4**). However, this improvement was less pronounced when considering complexes associated with the local network. Finally, at the level of the whole complexome, the behavior was essentially similar to the global dataset correlations. Our interpretation is that direct physical interactions within the local PPI network of selected gene(s) lend themselves to tight functional coherence that makes it much more likely for a higher mRNA-protein correlation. However, the scenario becomes significantly more complicated when considering larger groups (e.g., the complexome) since, especially in higher organisms, many proteins are shared subunits belonging to multiple complexes with high functional diversity among and/or between them.

## Discussion

Inborn errors of metabolism (IEMs) represent a group of genetic disorders that cause a disruption in the normal metabolic pathways of the cells, organs and organisms involved. These disorders manifest with a variety of symptoms, ranging from mild to life-threatening, and can cause long-term disability or even death. Understanding the precise molecular level changes in IEMs is key to developing clinical intervention strategies. Multiomics is a fast-growing field of study that combines data from multiple “omics” datasets (genomics, transcriptomics, proteomics, metabolomics, etc.) to generate a holistic picture of a disease state(s). By combining data from multiple omics datasets, multiomics provides a more comprehensive view of a disease, enabling researchers to gain deeper insights into the molecular pathways that underlie disease processes ^5,28,29^. This has particular relevance for the study of IEMs, as multiomics can provide a more detailed view of the molecular pathways involved in the etiology of these disorders. Functional studies to diagnose IEMs or confirm the relevance of identified genetic variants, often still occur via studies on skin-derived fibroblasts. There is a valid ongoing discussion whether these biospecimens are a suitable biological model to study the cellular disease mechanisms ^30^, but they remain a cornerstone of diagnostics and are therefore readily available for research on IEMs.

In this study, our analysis methodology exploited the systems-wide molecular landscape connectivity to identify how perturbations in single proteins (**Figure 1**) propagate across the system. In other words, do molecules that are significantly impacted function synergistically? Functional coherence is key to the coordinated organization of cellular pathways and molecular machines^31^. Analysis of omics (particularly proteomics) data by overlaying them together with annotated sets of protein complexes is an attractive approach^32^ as it provides clear insights into which processes and pathways are affected in the disease under study, IEMs in this work. Accordingly, based on the complexomics analysis, we found that multiple, functionally connected protein subunits that comprise the OXPHOS complexes were consistently affected (**Table 2** and **Figure 3B**). In addition to the OXPHOS structural proteins, a diverse group of additional mitochondrial complexes were also affected, involving practically every link in OXPHOS biogenesis: starting with mitochondrial DNA replication, mitochondrial transcription and translation via mitoribosomes to complexes regulating the import, assembly, synthesis and incorporation of mitochondrial complexes as well as mitochondrial morphology and cristae shaping (**Tables S1**). The complexomics framework also enabled cross-omics correlations to be made such as the link between changes in the composition of the lipid environment in Mito samples (**Figure 2**) which were then mapped to the relevant protein that functions in inter-organellar lipid transport involving the mitochondrion.

Starting from known information on the pathogenic variant in a gene linked to a disease (**Table 1**), the local interaction network analysis (**Figure 3A**) provides information about the direct interaction neighbourhood associated with the respective genetic mutation or the protein complex(es) directly involved with the mutated protein (**Table 2**). However, importantly, an enormous amount of information can be obtained from the global complexome analysis (see **Tables S1**). In-depth characterization of the proteome-level perturbation by analysing at the level of macromolecular protein assemblies (which represent the functional cellular machinery) elucidates the mechanisms underlying disease states, should lead to the identification of novel therapies and biomarkers and should allow us to get closer to understanding the phenotype of IEMs. Such a large-scale analysis of affected molecular machineries can also shed novel insights into and new implications for diseases such as IEMs. For example, the fact that chromatin remodelers and histone modifiers are affected in IEMs (**Figure 4**) represents a new lead. Therefore, the further detailed study of potential epigenetic dysfunction associated with metabolic disturbances can lead to a more novel characterization of IEMs, shed light on hitherto unexplained phenotypical traits, thereby leading to a better mechanistic understanding and potentially novel treatment strategies.

The complexome approach outlined here is a generally applicable methodology that can also be used using transcriptomics data as a proxy (if corresponding proteomics data is unavailable). Complexomes are currently available for all major model organisms, so the method should be widely applicable. In addition to Complex Portal (used here), there are a number of other resources (e.g., CORUM^33^ and hu.MAP 2.0^34^) where the focus is also on curating protein assemblies with clearly defined cellular functions. These data intensive resources are also regularly updated, with the numbers of annotated complexes as well as the organismal coverage increasing consistently^35^. In summary, this type of analysis protocol described here should be widely applicable to diverse disease relevant biological investigations, that will allow researchers to prioritize target complexes/pathways for further detailed characterization and inform potential therapy options for clinical use.

## Materials and Methods

### Samples

Fibroblasts from patients with different genetically confirmed IEM and control fibroblasts cell lines obtained from healthy individuals were retrospectively recruited from the UZ Leuven fibroblast databank. We included in this study 5 control cell lines, 6 cell lines derived from individuals with the organic acidaemia methylmalonic aciduria (3 *MMAB* defects and 3 *CblC* defects), 4 cell lines with a beta-oxidation defect (2 very long-chain acyl-CoA dehydrogenase deficiency and 2 medium-chain acyl-CoA deficiency) and 6 cell lines with an OXPHOS deficiency (Complex I deficiency). The fibroblasts cell lines were maintained in DMEM medium with 5.5 mM glucose (closest approximation to physiological glycemia) and 2 mM glutamine supplemented with 10% foetal bovine serum at 37 °C with 5% CO2 in a humidified incubator. Cells were not allowed to grow further than passage 15. Cells were screened for Mycoplasma infection (Westburg).

Fibroblasts were plated at 2,500 cells/cm^2^/0.15 mL of DMEM (ThermoFisher) with 5.5 mM Glucose, 2 mM Glutamine. Samples for the transcriptomics analysis were plated in T175 culture flasks, samples for the metabolomics and proteomics analyses were plated in 6 well plates. At 72 h, samples were collected by collecting the media and harvesting the cells by scraping on ice. All samples were prepared in triplicates.

The analyses of fibroblasts were approved by the ethical committee of the University Hospitals of Leuven under application number S60206 (retrospective metabolic analysis of archived fibroblasts) and S58977 (prospective assessment of mitochondrial function in fibroblasts). Informed consent was obtained from the patients or their legal guardian(s).

### Proteomics Sample preparation

First, protein pellets were dried in a vacuum concentrator to remove all residual methanol and denatured in a buffer containing 8 M urea and 20 mM HEPES. Proteins were reduced by addition of 15 mM dithiothreitol (DTT) and incubation for 30 minutes at 55°C and then alkylated by addition of 30 mM iodoacetamide (IAA) for 15 minutes at room temperature in the dark. Samples were diluted with 20 mM HEPES pH 8.0 to a final urea concentration of 4 M and proteins were digested with 1 µg lysyl endopeptidase (Wako) (1/100, w/w) for 4 hours at 37°C. Samples were again diluted to 2 M urea and digested with 1 µg trypsin (Promega) (1/100, w/w) overnight at 37°C. The resulting peptide mixture was acidified by addition of 1% trifluoroacetic acid (TFA) and desalted on a reversed phase (RP) C18 OMIX tip (Agilent). The tip was first washed 3 times with 200 µl pre-wash buffer (0.1% TFA in water/acetonitrile (ACN, 20:80, v/v)) and pre-equilibrated 5 times with 200 µl of wash buffer (0.1% TFA in water) before the sample was loaded on the tip. After peptide binding, the tip was washed 3 times with 200 µl of wash buffer and peptides were eluted twice with 100 µl elution buffer (0.1% TFA in water/ACN (40:60, v/v)). The combined elutions were dried in a vacuum concentrator.

### LC-MS/MS analysis

Purified peptides were re-dissolved in 20 µl loading solvent A (0.1% TFA in water/ACN (98:2, v/v), the peptide concentration was determined on a Lunatic^36^ spectrophotometer (Unchained Labs) and 2 µg of each sample was injected for LC-MS/MS analysis on an Ultimate 3000 RSLCnano system in-line connected to an Orbitrap Fusion Lumos mass spectrometer (Thermo Scientific) equipped with a pneu-Nimbus dual ion source (Phoenix S&T). Trapping was performed at 10 μl/min for 4 min in loading solvent A on a 20 mm trapping column (made in-house, 100 μm inner diameter(ID), 5 μm beads, C18 Reprosil-HD, Dr. Maisch) and the sample was loaded on a 200 cm long micro pillar array column (Thermo Scientific) with C18-endcapped functionality mounted in the Ultimate 3000’s column oven at 50°C. For proper ionization, a fused silica PicoTip emitter (10 µm ID, New Objective) was connected to the µPAC™ outlet union and a grounded connection was provided to this union. Peptides were eluted by a non-linear increase from 5 to 55% MS solvent B (0.1% FA in water/ACN (2:8, v/v)) over 145 minutes, first at a flow rate of 750 nl/min, then at 300 nl/min, followed by a 15-minutes wash reaching 99% MS solvent B and re-equilibration with 95% MS solvent A (0.1% FA in water).

The mass spectrometer was operated in data-dependent mode, automatically switching between MS and MS/MS acquisition. Full-scan MS spectra (300-1500 m/z) were acquired in 3 s acquisition cycles at a resolution of 120,000 in the Orbitrap analyzer after accumulation to a target AGC value of 2E5 with a maximum injection time of 250 ms. Monoisotopic precursor selection (MIPS) was set to Peptide and the precursor ions were filtered for charge states (2-7 required), dynamic exclusion (60 s; +/- 10 ppm window) and intensity (minimal intensity of 5E3). The precursor ions were selected in the multipole with an isolation window of 1.2 Da and accumulated to an AGC target of 1.2E4 or a maximum injection time of 40 ms and activated using HCD fragmentation (34% NCE). The fragments were analysed in the Ion trap Analyzer with Normal scan rate. The polydimethylcyclosiloxane background ion at 445.120028 Da was used for internal calibration (lock mass). QCloud^37^ was used to control instrument longitudinal performance during the project.

### Proteomics data analysis and identification of differentially expressed (regulated) proteins

LC-MS/MS runs of all 62 samples were searched together using the MaxQuant algorithm (version 1.6.2.6) with mainly default search settings, including a false discovery rate set at 1% on peptide and protein level. Spectra were searched against the human reference proteome sequences in the Uniprot database (database release version of 2020_01), containing 20.595 sequences (https://www.uniprot.org). The mass tolerance for precursor and fragment ions was set to 20 and 4.5 ppm, respectively, during the main search. Enzyme specificity was set to C-terminal of arginine and lysine, also allowing cleavage at Arg/Lys-Pro bonds with a maximum of two missed cleavages. Variable modifications were set to oxidation of methionine residues and acetylation of protein N-termini whereas carbamidomethylation of cysteine residues was set as fixed modification. In order to reduce the number of spectra that suffer from co-fragmentation, the precursor ion fraction (PIF) option was set to 75%. Only proteins with at least one unique or razor peptide were retained leading to the identification of 4,749 proteins Proteins and peptides were quantified by the MaxLFQ algorithm, integrated in the MaxQuant software. A minimum ratio count of two unique or razor peptides was required for quantification.

Next, the proteinGroups.txt file containing the aggregated protein intensities obtained from MaxQuant was processed using the PaDuA python package^38^ that is optimized to work with MaxQuant output data and follows closely typical Perseus workflows. Reverse database hits, potential contaminants and proteins only identified by site were first removed. Then LFQ intensities were log2 transformed and replicate samples were grouped. Proteins with valid values in at least 2 out of 3 replicates from at least one patient sample were retained (3140 proteins in total). Missing values were then imputed from a normal distribution with default width of 0.3 and down shift of 1.8. Column-wise imputation was applied, where imputation is applied to each expression column separately. To compare protein intensities in different sample groups, statistical testing for differences between the two-group means was performed, using two-sample independent t-tests.

### Transcriptomics experimental setup

RNA concentration and purity were determined spectrophotometrically using the Nanodrop ND-8000 (Nanodrop Technologies) and RNA integrity and concentration were assessed using a Fragment Analyzer SS-RNA kit (Agilent). Per sample, an amount of 500 ng of total RNA was used as input. Using the Illumina TruSeq® Stranded Total RNA Sample Prep Kit with Ribo-Zero Gold (protocol version “1000000040499 v00 - October 2017”) rRNA is depleted from the total RNA samples using Ribo-Zero Gold ribosomal RNA reduction chemistry. Subsequently, RNA was purified and fragmented to a limited extent (allowing for longer RNA fragments) and converted into first strand cDNA in a reverse transcription reaction using random primers. Next, double-stranded cDNA was generated in a second strand cDNA synthesis reaction using DNA PolymeraseI and RNAse H. The cDNA fragments were extended with a single ‘A’ base to the 3’ ends of the blunt-ended cDNA fragments after which multiple indexing adapters were ligated introducing different barcodes for each sample. Finally an enrichment PCR was carried out to enrich those DNA fragments that have adapter molecules on both ends and to amplify the amount of DNA in the library. Sequence-libraries of each sample were equimolar pooled and sequenced on Illumina NovaSeq 6000 (v1.5 kit, S1 flowcell), paired end reads 150 (151-8-8-151) at the VIB Nucleomics Core (https://nucleomicscore.sites.vib.be/).

### Transcriptomics data processing

#### Read preprocessing

FastX 0.0.14 (HannonLab. Fastx-toolkit. 2010) was used to trim low quality ends (< Q20). Reads shorter than 35bp after trimming were removed. Cutadapt 1.15 ^39^ was then used to perform adapter trimming. The adapters were trimmed only at the end (at least 10bp overlap and 90% match). Reads that were shorter than 35bp after adapter trimming were removed. Next using FastX 0.0.14 and ShortRead 1.40.0 ^40^, we performed a quality filtering by removing polyA-reads (more than 90% of the bases equal A), ambiguous reads (containing N), low quality reads (more than 50% of the bases < Q25) and artifact reads (all but 3 bases in the read equal one base type). Finally, since the filtering of reads may remove one read of a pair and make paired fastq-files inconsistent, pairing was made consistent by removing reads that belong to broken pairs.

### Mapping

We aligned the preprocessed reads to the reference genome of Homo sapiens Ensembl.GRCh38.88 (GRCh38) using STAR 2.5.2b ^41^. Default STAR aligner parameter settings were used, except for the following: “--outSAMprimaryFlag OneBestScore --twopassMode Basic --alignIntronMin 50 --alignIntronMax 500000 –alignMatesGapMax 500000 --outSAMtype BAM SortedByCoordinate”. Next using samtools 1.5, we removed reads from the alignment that were non-primary mappings or had a mapping quality <= 20 ^42^. Also with samtools 1.5 we sorted the reads from the alignment according to the chromosomes and indexed the resulting bam-files.

### Counting

We counted the number of reads in the alignments that overlap with the gene features using featureCounts 1.5.3 ^43^. We chose the following parameters: -pBC -Q 0 -s 2 -t exon -g gene_id. Reads that can be attributed to more than one gene (ambiguous) or could not be attributed to any gene (no feature) were not counted. We also removed genes for which all samples had less than 1 counts per million. We further corrected raw counts within samples for GC-content and between samples using full quantile normalization, with the EDASeq package from Bioconductor ^44^.

### Differential gene expression analysis

EdgeR 3.38.4 package of Bioconductor was used to fit a negative binomial generalized linear model (GLM) to the count data ^45^. Differential expression was tested with a GLM likelihood ratio test, also implemented in EdgeR. The resulting p-values were corrected for multiple testing with Benjamini-Hochberg to control the false discovery rate (FDR).

### Untargeted metabolomics experimental setup

Prior to metabolite extraction, the cells were washed with a cold (4°C) 0.9% NaCl solution. After removal of the wash solution, 300µL 80% cold MeOH was added to the cells. After 3 minutes of incubation on ice, the cells were scraped and the extraction mix was transferred to a new Eppendorf. Subsequently, the samples were centrifuged for 15 min at 4°C using 20,000xg. 250µL of the MeOH extract was transferred to a new Eppendorf and evaporated to dryness under vacuum conditions. The dried extracts were resolubilized in 40 µL 80% MeOH. Samples were subjected to Ultra Performance Liquid Chromatography High Resolution Mass Spectrometry (UPLC-HRMS) at the VIB Metabolomics Core Ghent (VIB-MCG). 10 ul was injected on an Acquity UHPLC (Waters) device connected to a Vion HDMS Q-TOF mass spectrometer (Waters). All biological samples were analyzed at random and pooled samples were included for system reproducibility. Chromatographic separation was carried out on an ACQUITY UPLC BEH C18 (50 x 2.1 mm; 1.7 μm) column (Waters) with the column temperature maintained at 40 °C. A gradient of two buffers was used for separation: buffer A (99% water, 1% acetonitrile and 0.1% formic acid, pH 3) and buffer B (99% acetonitrile, 1% water and 0.1% formic acid, pH 3), as follows: 99 % A decreased to 50% A in 5 min, decreased to 30% from 5 to 7 minutes, and decreased to 0% from 7 to 10 minutes. The flow rate was set to 0.5 mL/min. Electrospray Ionization (ESI) was applied, with the LockSpray ion source operated in negative and positive ionization mode under the following specific conditions: capillary voltage, 2.5 kV; reference capillary voltage, 3 kV; source temperature, 120°C; desolvation gas temperature, 600°C; desolvation gas flow, 1000 L/h; cone voltage, 40 V and cone gas flow, 50 L/h. The collision energy for full MS scan was set at 4 eV for low energy settings, for high energy settings (HDMSe) it was ramped from 20 to 70 eV. Mass range was set from 50 to 1200 Da and scan time was set at 0.1s. Nitrogen (greater than 99.5%) was used as desolvation and cone gas. Leucine-enkephalin was used for the lock mass calibration. Profile data was recorded through Unifi Workstation v2.0 (Waters) and data processing was performed with Progenesis QI software version 3.0 (Waters) for chromatogram alignment and compound ion detection. The detection limit was set at maximum sensitivity and the data was normalized to the protein weight. In ESI- and ESI+ ionization, 1,866 and 4,889 compound ions were detected respectively and the chromatograms were aligned to “PooledSample_02”. Statistical analyses (ANOVA p-value < 0.05) were performed on ArcSinh-transformed and pareto scaled ion intensities. For pairwise comparison a p-value of 0.05 and fold change of 2 were applied. Compound annotation for significant features was done by using MS-FINDER v3.57 ^46,47^. Massbank, GNPS, ReSpect and an in house database, PhytoFrag were used for spectral matching. The following databases were used for formula prediction and structural matching by in silico fragmenter: HMDB, Urine, Saliva, Feces, Serum, CSF, SMPDB, LipidMAPS, YMDB, BMDB , NPA, and an in house database, PhytoComp. The following parameter settings were applied: precursor mass tolerance: 5 ppm, fragment mass tolerance: 50 ppm, and element selection: C, H, N, O, P and S.

### Protein extraction and immunoblotting

Fibroblasts were cultured in low glucose Dulbecco’s Modified Eagle’s Medium (DMEM) (Gibco™, ref. 31885049) supplemented with 10% fetal bovine serum (FBS), in a humidified incubator at 37°C with 5% CO_2_. Once they reached a 80-90% confluence, the fibroblasts were scraped in PBS, and histones were extracted using a histone extraction kit (Abcam, Cat# ab113476), according to the manufacturer’s instructions. Protein concentration was measured in the supernatant using the BCA protein assay kit (Thermo Scientific, Cat# 23225). Samples were separated on a 10% Bis-Tris gel (Invitrogen, Cat# NP0315BOX) under reducing conditions, transferred to a nitrocellulose membrane (Invitrogen, Cat# LC2001), and analyzed by immunoblotting. Mouse monoclonal pan acetylation antibody (Proteintech, Cat# 66289-1-Ig) was used at a 1:1000 dilution, and rabbit β-Actin antibody (Cell Signaling, Cat# 4967) at 1:2000. Anti-mouse (Cell Signaling, Cat# 7076) and anti-rabbit (Cell Signaling, Cat# 7074) HRP-linked secondary antibodies were used at 1:2000. Signal was detected using the ECL Western Blotting Substrate (Thermo Scientific, Cat# 32209) according to the manufacturer’s instructions. Densitometric quantification of the proteins in the bands was performed using the Fiji software.

### GO Biological Process over-representation statistical testing based on lists of regulated proteins/transcripts

PANTHER^15^ was used to perform the statistical over-representation tests. This test allowed us to identify statistically over-represented biological processes among the list of regulated proteins (compared to the entire human proteome as background reference) for each individual disease category (OA, FA and Mito). We used both ‘Complete’ and ‘Slim’ GO Biological Process annotation sets (Release dated 20220712). The former includes both manually curated and electronically generated GO annotations, whereas the latter is based on manually curated annotations alone. GO Ontology database (Released 2022-07-01)^48^.

## Supplementary Figure Legends

**Figure S1.**
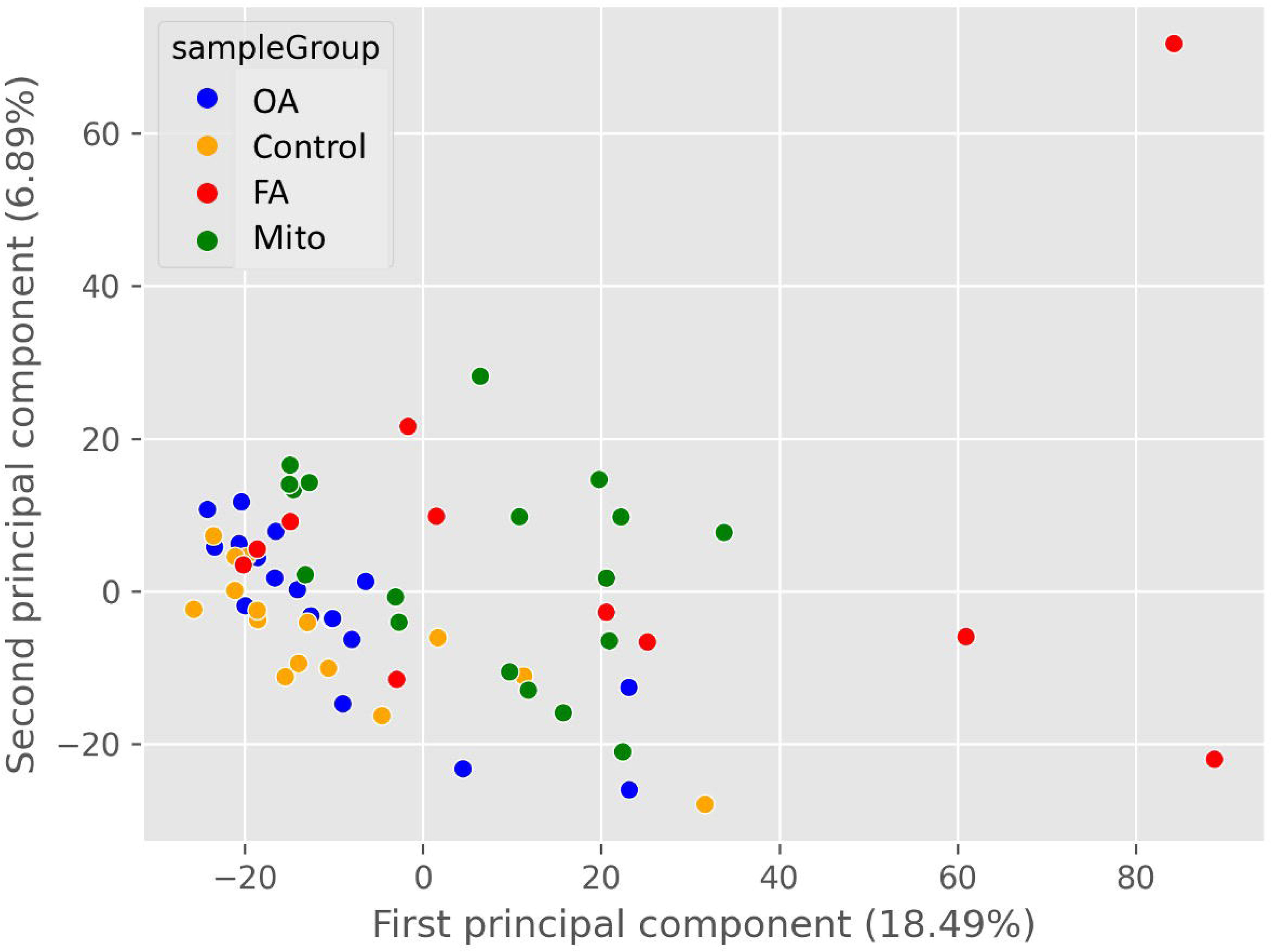
Principal Component Analysis (PCA) plot based on the proteomics data. IEM groups are shown as indicated by the different colours. Plotted using the Python-based PaDuA pipeline ^38^.

**Figure S2.**
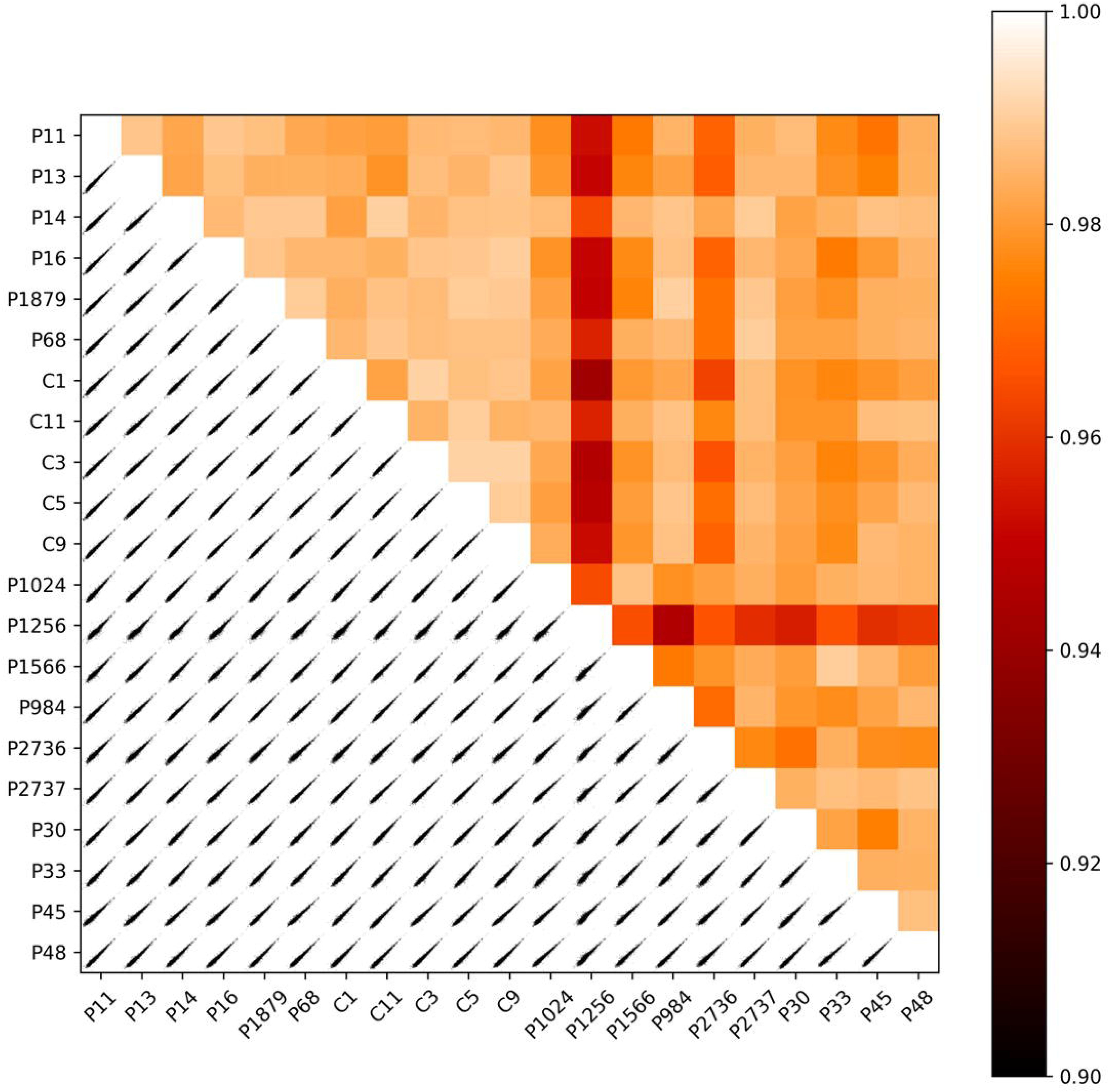
Heatmap of the Pearson correlation coefficients (color bar is shown) based on the protoemics data, calculated pairwise between the IEM samples. Sample names are indicated. Bottom half of the plot shows the equivalent scatter plots of the protein intensities measured in the corresponding datasets. Plotted using the Python-based PaDuA pipeline ^38^.

**Figure S3.**
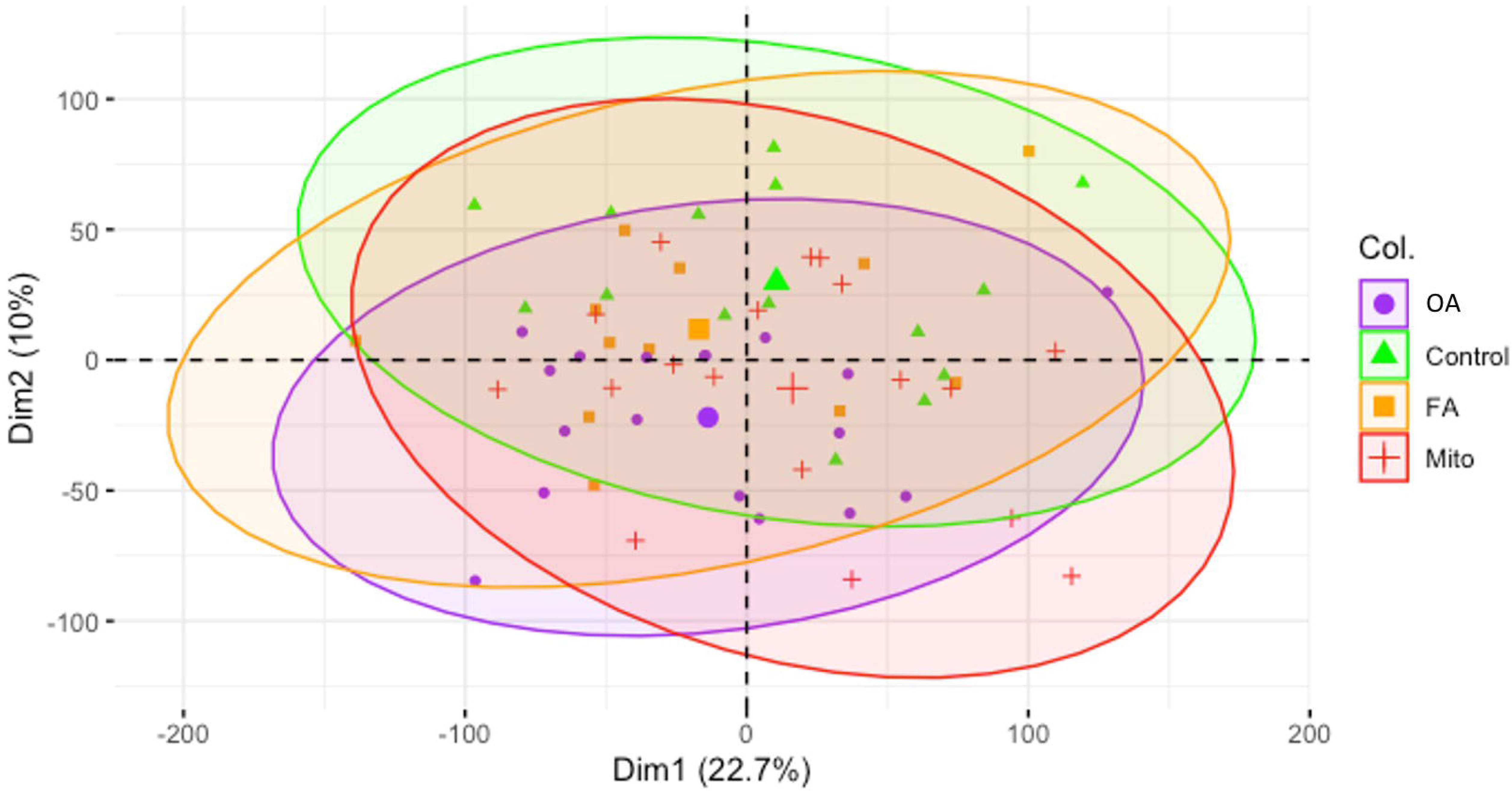
Principal Component Analysis (PCA) plot based on the transcriptomics data. IEM groups are shown as indicated by the different colours. Plotted using R (https://www.R-project.org/).

**Figure S4.**
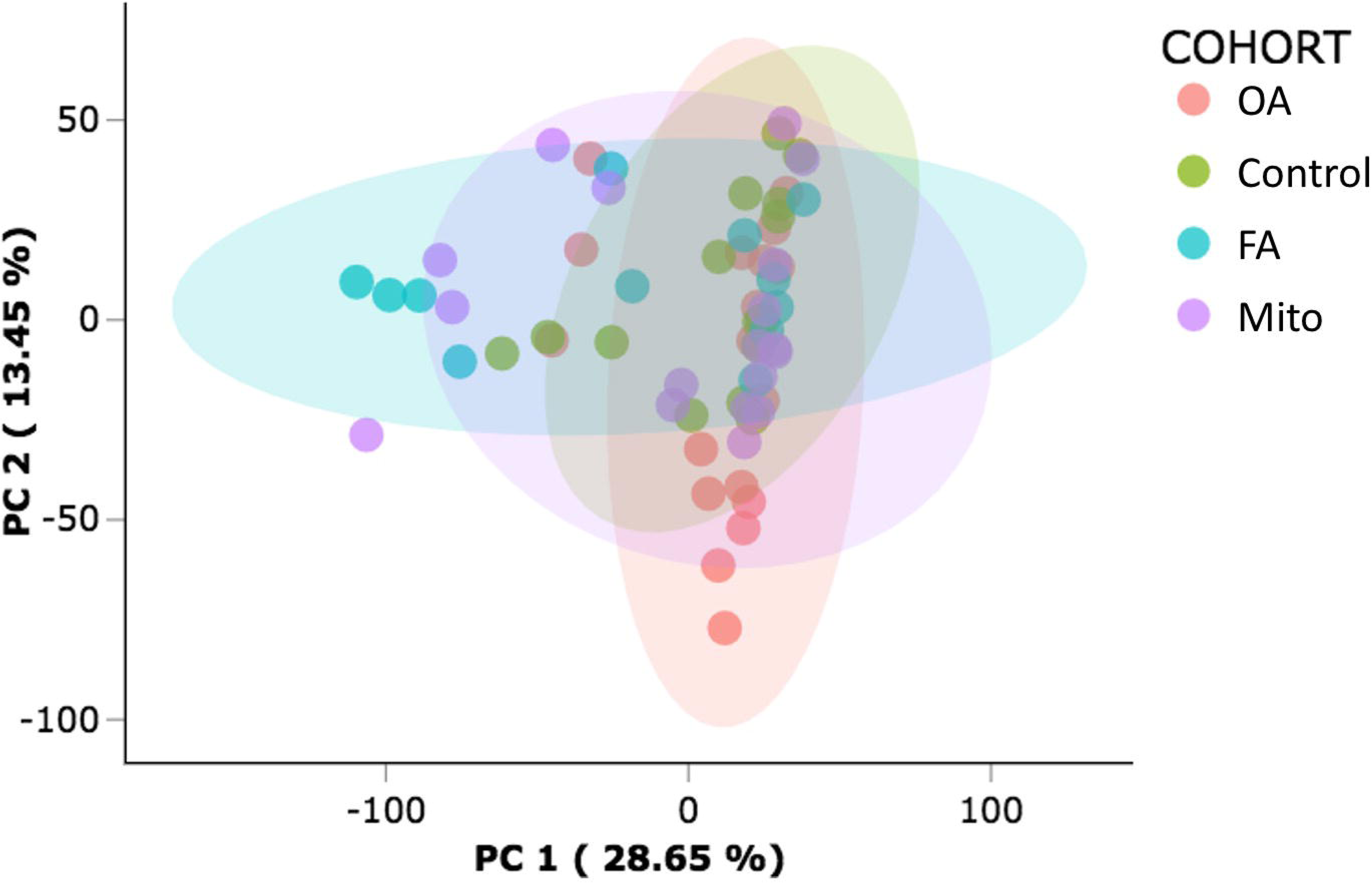
Principal Component Analysis (PCA) plot based on the untargeted metabolomics data. IEM groups are shown as indicated by the different colours. Plotted using an Elucidata^TM^ (https://www.elucidata.io) metabolomics data processing pipeline provided by the VIB Metabolomics Core, Ghent.

**Figure S5.**
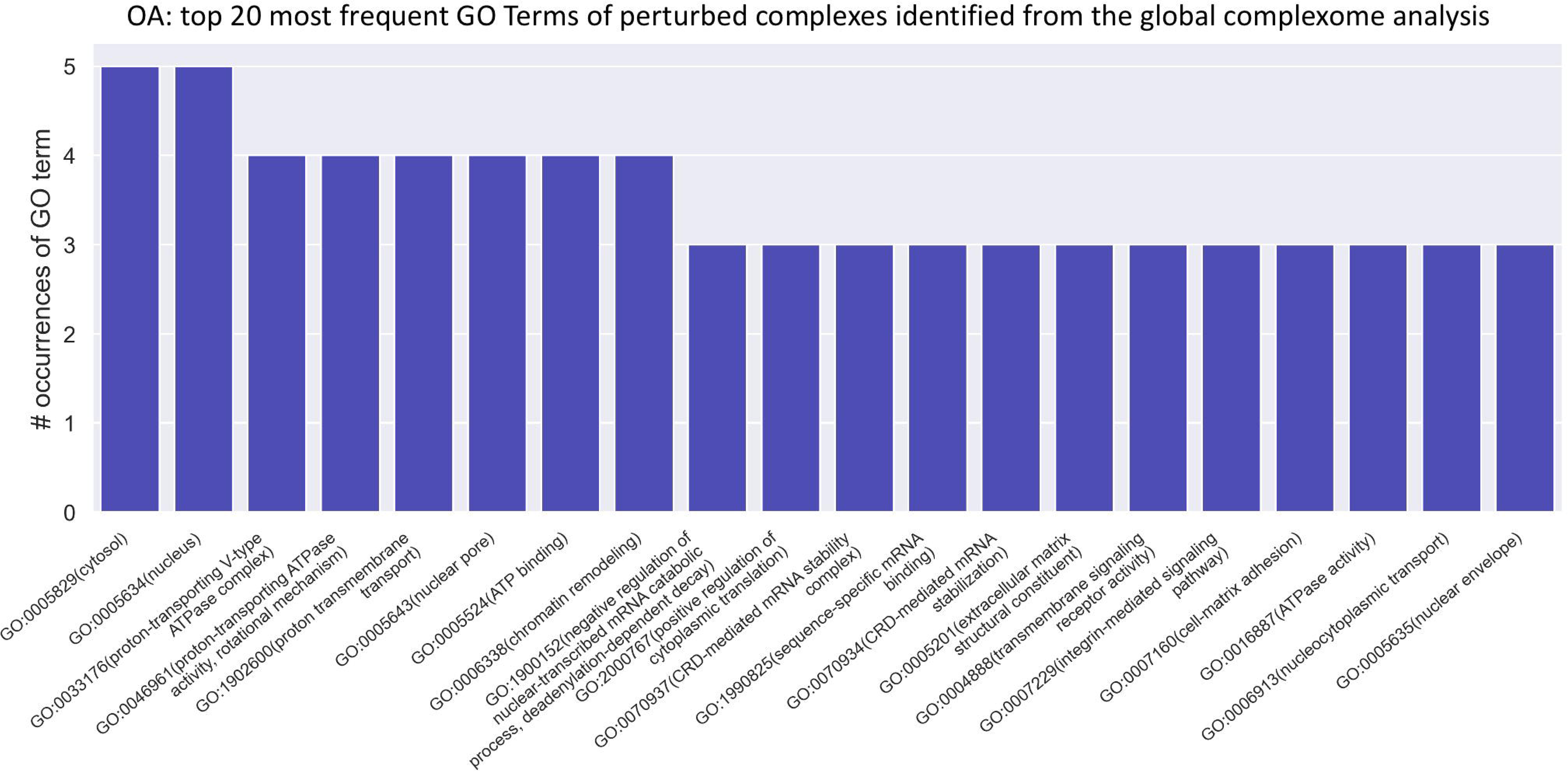
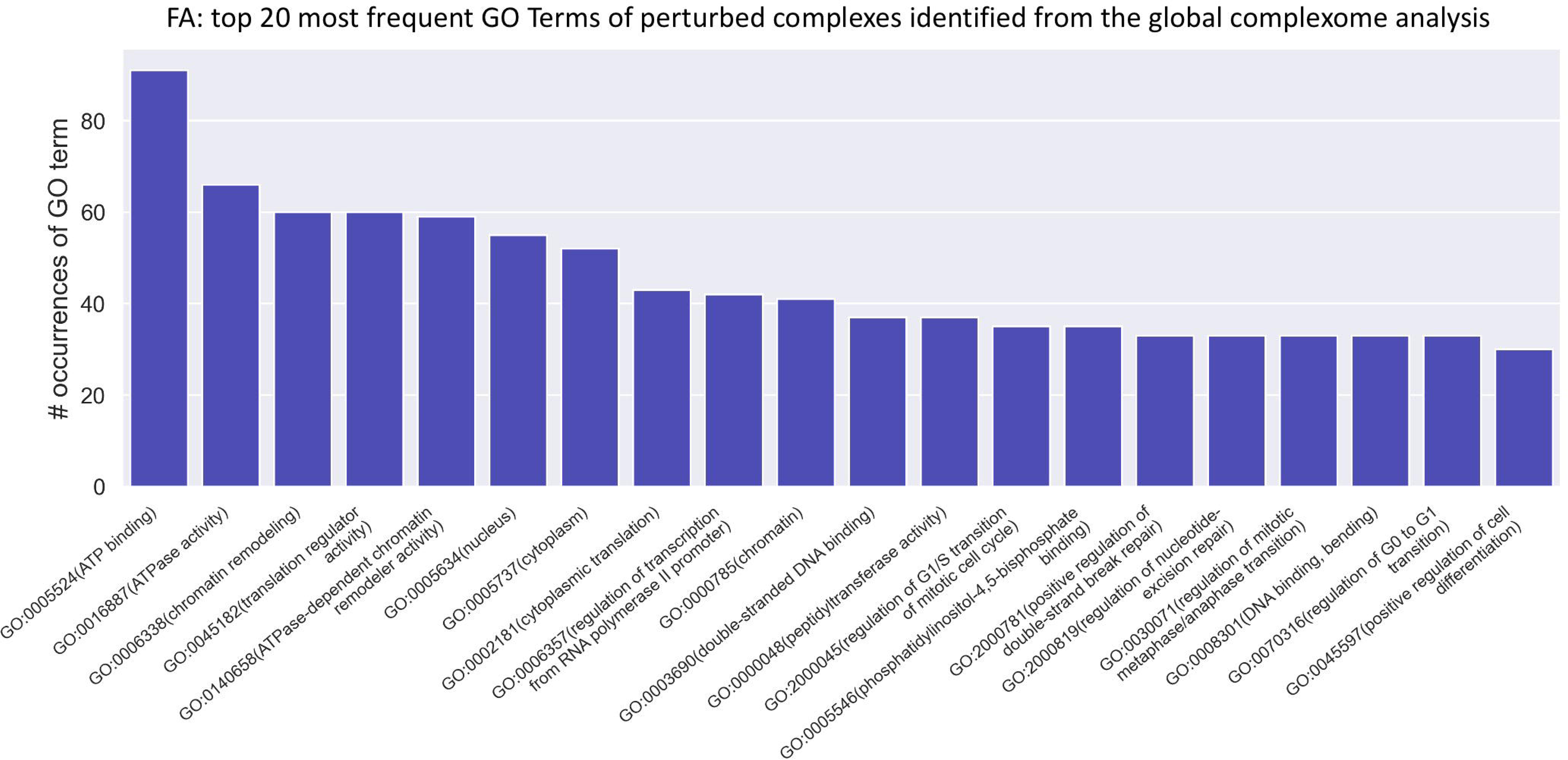
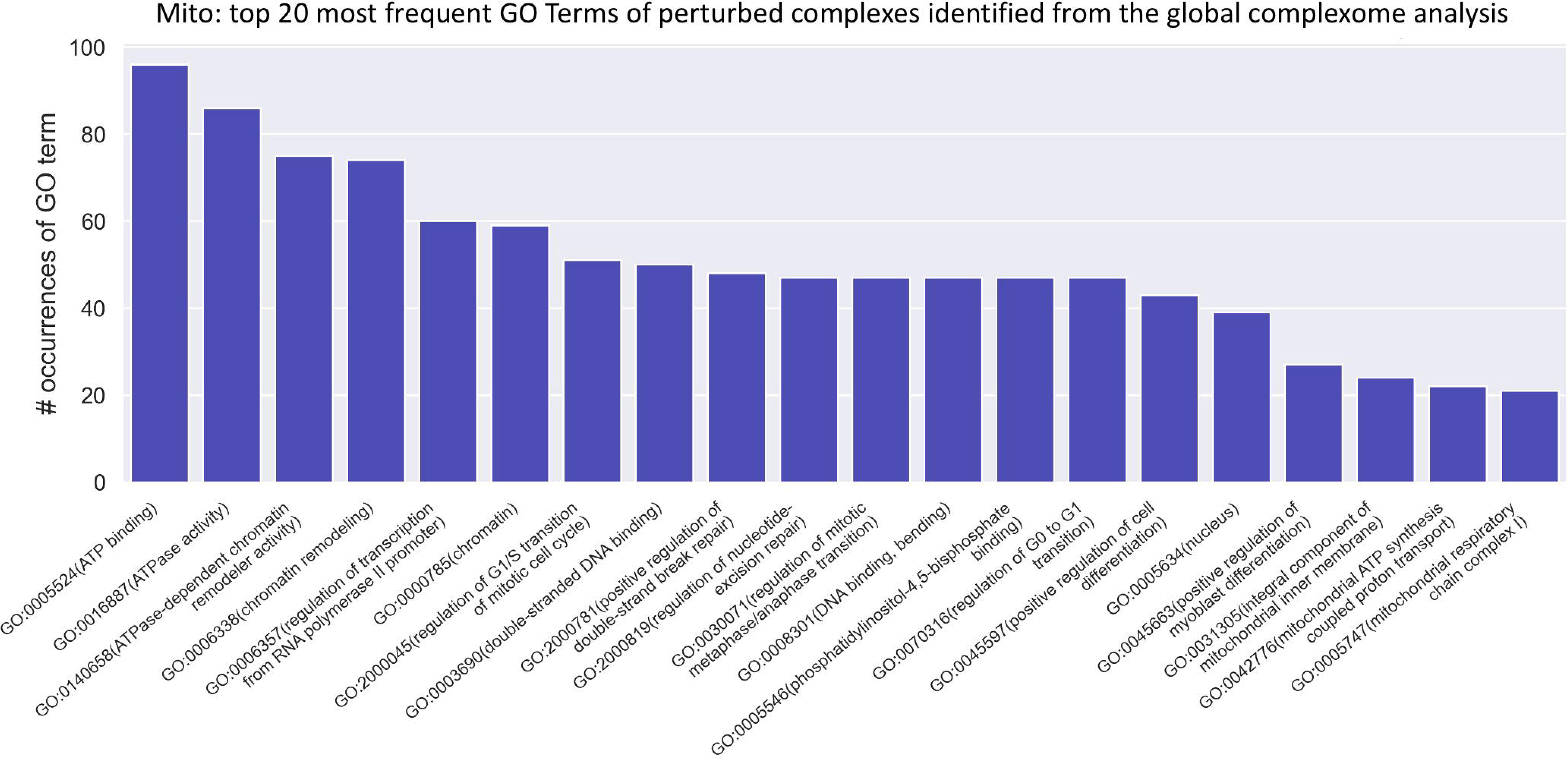
Plot per IEM category, showing the top 20 most frequently annotated GO terms mapped to the perturbed protein complexes (identified based on the global complexome analysis) within each IEM category.

## Acknowledgements

This work was supported by the Katholieke Universiteit Leuven (C1 grant: EFF-D2860-C14/17/110) (DC, BG) and the Tjallingh Roorda Foundation (IA). PW was funded by the Fonds Wetenschappelijk Onderzoek-Vlaanderen (Fundamenteel Klinisch Mandaat 18B4322N). The authors wish to thank Christof De Bo (VIB, Ghent) for help with drawing Figure 3.

## Notes

### Competing Interest Statement

The authors have declared no competing interest.

